# A 3T investigation of *B*_1_^+^-inhomogeneity tolerance in MP2RAGE-based *R*_1_-mapping by calculating second contrast that permits previously problematic sequence parameters

**DOI:** 10.1101/2024.04.10.588855

**Authors:** Lenno Ruijters, Torben Ellegaard Lund, Mads Sloth Vinding

**Author notes:** Corresponding author: Lenno Ruijters.

## Abstract

**Purpose:** Regarding in vivo, robust *R*_1_-mapping, the goal of the present paper is twofold. First, to verify that non-bijective mapping in MP2RAGE imaging can be resolved through a 2D look-up-table approach. Second, that the expanded parameter space from this can be used to improve *B*_1_^+^-inhomogeneity tolerance without other prerequisites.

**Theory:** By deriving a second contrast from the magnitude images of the MP2RAGE acquisition, ambiguities in the original MP2RAGE image resulting from non-bijective transfer curves can be resolved. Such ambiguities may occur when protocols are optimised, e.g., for higher *B*_1_^+^-inhomogeneity tolerance. A 2D look-up-table approach combines the available information to resolve these ambiguities during mapping.

**Methods:** At 3T, we acquired MP2RAGE images with standard acquisition parameters and (non-bijective) parameters optimised for *B*_1_^+^-inhomogeneity tolerance. From three subjects across multiple sessions, we assessed the *B*_1_^+^-inhomogeneity tolerance through excitation pulse amplitude scalings.

**Results:** The *R*_1_-maps resulting from the *B*_1_^+^-optimised protocols showed greatly reduced *B*_1_^+^-effects across images, but without additional scanner time. Meanwhile these maps could only successfully be derived by a 2D look-up-table approach.

**Conclusion:** We show that it is possible to optimise for *B*_1_^+^-inhomogeneity tolerance in MP2RAGE through sequence parameter settings, while still successfully estimating the *R*_1_-map with a 2D look-up-table approach. This without the need for an additional *B*_1_^+^-map. The increased parameter space enabled by the 2D look-up-table approach may further be used to adjust MP2RAGE acquisitions for improved scan times, SNR and/or CNR.

## 1. Introduction

In order for *R*_1_-mapping to achieve subject-specific, in-vivo, detailed maps of human cortical structures (building on Refs.^1–3^), it is important that *R*_1_-changes reflect variations in myelo-architecture rather than contaminations from artefacts. To map structurally distinct cortical areas beyond primary sensory tissue^1–3^, i.e. whole-brain, highly refined images are required. In order to detect subtle variations, these images need to be detailed and reliable, i.e. have high resolution and minimal artefact contributions. One prominent source of artefacts is *B*_1_^+^-inhomogeneity, which modulates the flip angle (FA), and consequently the measured signal. Factors that affect the size of *B*_1_^+^-effects include *B*_0_, readout train length, and *T*_R_. The result is that, as higher resolutions become possible, *B*_1_ ^+^-artefacts increasingly counteract the benefits of increased spatial detail. Methodological strategies are necessary to limit the loss in *R*_1_-specificity as increased resolutions are achieved.

MP2RAGE is a *T*_1_-weighted MRI-sequence that produces quantitative maps, while simultaneously mitigating *B*_1_-effects^4^. By acquiring two Rapid Gradient Echoes, and by combining these during post-processing into a unified MP2RAGE image, *T*_2_*, *B*_0_ and *B*_1_^-^ effects can be eliminated. Calibrating sequence parameters, like the FAs or inversion times, can further minimise the *B*_1_ ^+^-effects. Novel pTx additionally addresses *B*_1_ ^+^-effects^5–9^, although accessible implementations leave residual *B*_1_^+^-inhomogeneities^8^. Such residual inhomogeneities will cause issues with, e.g., increased number of slices.

Since MP2RAGE images are quantitative, the image values can be derived from *T*_1_ and the sequence parameters via Bloch simulations. If the mapping is bijective (i.e. one-to-one), then each MP2RAGE value can be uniquely associated with a *T*_1_-value^10^. However, parameters with desirable features, including improved tolerance to *B*_1_^+^-inhomogeneities, but also high within- or between-tissue contrast, or faster scanning times, may produce mappings that are non-bijective. Consequently, *R*_1_-values are misestimated and a different strategy is required to successfully derive *R*_1_-maps.

In this paper, we investigate the possibility of mitigating *B*_1_^+^-effects with modified sequence parameters, by tackling the challenge of non-bijective mappings between MP2RAGE values and *R*_1_. To this end, we introduce a new strategy able to derive *R*_1_-maps from MP2RAGE acquisitions with non-bijective mappings. The present *R*_1_-mapping utilises a secondary contrast, derived from the MP2RAGE acquisition, based on the ratio between the difference and the sum of the magnitude of the two gradient echo images (Difference-Sum-Ratio; DSR).

Similar to the MP2RAGE image, effects related to *T*_2_*, *B*_0_ and *B*_1_^-^ are cancelled out in the DSR image. Additionally, the DSR-contrast is suitable for disambiguation of MP2RAGE values. Consequently, accurate *R*_1_-maps can be determined for non-bijective mappings, as may be the case in protocols highly tolerant of *B*_1_^+^-inhomogeneity.

## 2. Theory

Quantitative *T*_1_-weighted images are derived from the MP2RAGE acquisition by combining the gradient echo images (*I*_1_, *I*_2_), acquired from each inversion time (*T*_I,1_, *T*_1, 2_), voxel by voxel into a unified image (*I*_*UNI*_):

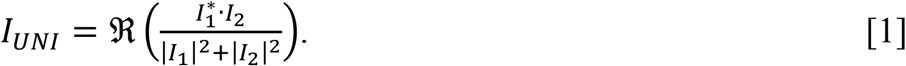

When nuisance factors contribute equally to *I*_*1*_ and *I*_*2*_, these effects cancel out in *I*_*UNI*_. By nature of Eq. (1), *I*_*UNI*_-values are bound between -0.5 and +0.5, and exactly equal to ±0.5, when *I*_*1*_ = ±*I*_*2*_. Because these values are quantitative, *I*_*UNI*_-values can be derived from *T*_1_ or *R*_*1*_(=1/*T*_*1*_). In standard implementations, *I*_*UNI*_ is mapped to *R*_1_-values via a 1-dimensional look-up table (1D-LUT), which can be visualised by a transfer curve (see Fig 1). A bijective relationship between *I*_*UNI*_ and *R*_1_ is required for the 1D-LUT procedure to be applicable. This is achieved by careful selection of sequence parameters (Fig 1a)^4^. By scaling the FAs, the effects of *B*_1_^+^-inhomogeneity on estimated *R*_1_-values can be derived and visualised as separate transfer curves.

**Figure 1:**
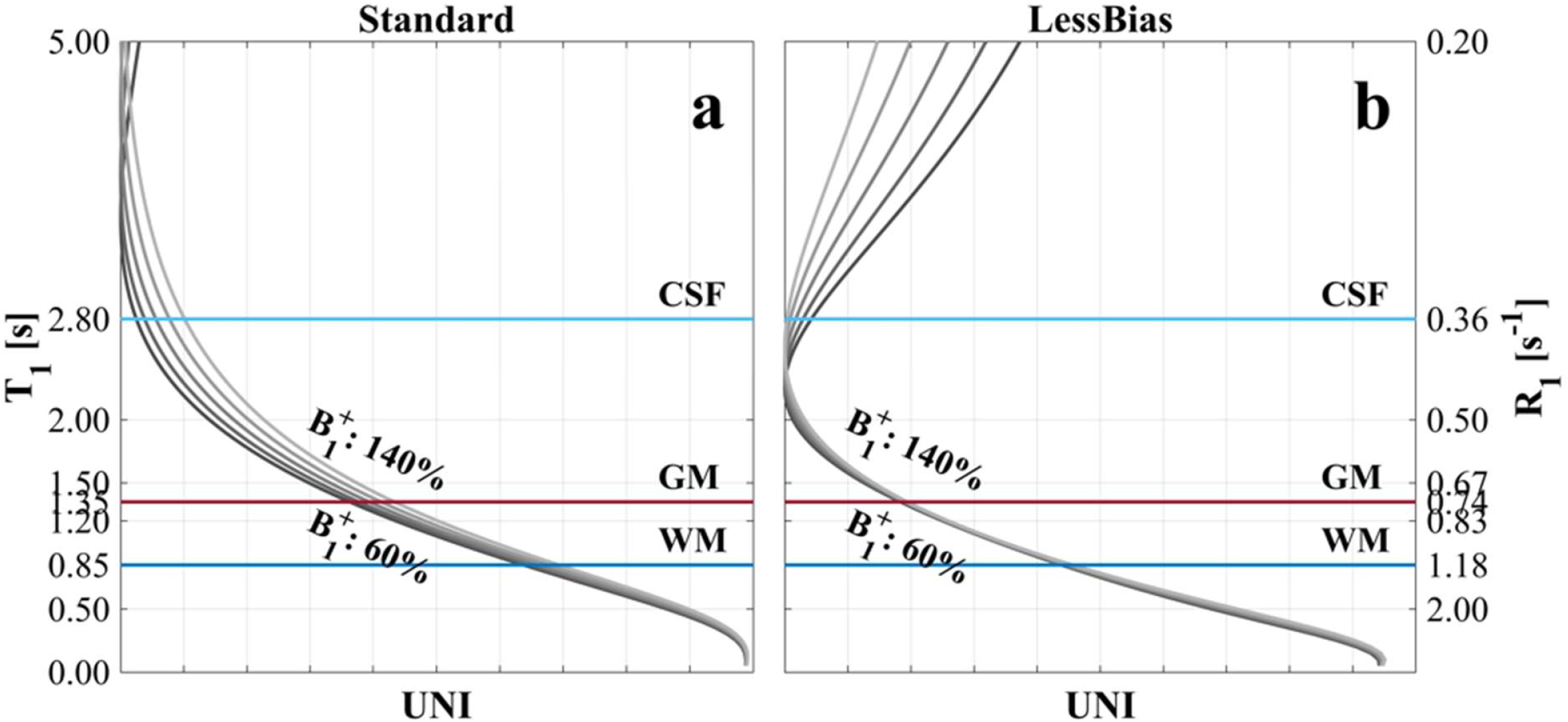
The MP2RAGE-to-T_1_ transfer curves for the Standard (a) and LessBias (b) protocols, with corresponding R_1_-values on the right axis. The Standard protocol, (a), is more sensitive to B ^+^-inhomogeneities than the LessBias protocol, (b), which is almost insensitive to B ^+^-variations in the order of +/-40%. Within the range of T -values from 0 to 5 s the Standard protocol, (a), has a near-bijective relationship between the MP2RAGE UNI values and T_1_; this is not the case for the LessBias protocol, (b), where within the T_1_-window, there are ambiguous UNI values. Also notice we maintain a near identical contrast between GM and WM.

It is possible to select sequence parameters where the corresponding transfer curve results in less dependency on *B*_1_^+^-variations (see Fig 1b). However, such parameters may lead to non-bijective mapping within the *R*_1_-window-of-interest. Here, the 1D-LUT method fails, because *R*_1_-values from two different tissue types (e.g. grey matter (GM) and cerebrospinal fluid (CSF)) map to the same *I*_*UNI*_-value. These non-bijections will occur when *I*_*1*_=±*I*_*2*_ at any point within the *R*_1_-window-of-interest. For standard protocols, non-bijection will typically occur for low *R*_1_s (≈<0.2s^-1^), leading to large *R*_1_-variance within CSF, which is normally unproblematic (however, see Discussion). For mappings where the non-bijection occurs at relatively high *R*_1_-values (like in Fig 1b), a solution is needed (see Supporting Information, section 3).

By introducing a second, distinct contrast that is derived from *I*_*1*_ and *I*_*2*_, we are able to solve ambiguities in non-bijective mappings, and accurately estimate the *R*_1_-values. This second contrast can be defined in several ways. We settled on a combination of the magnitudes of *I*_*1*_ and *I*_*2*_, since these magnitude images are readily available in standard setups. Similar to *I*_*UNI*_, we further wanted this contrast to be quantitative and tolerant to magnetic field inhomogeneities. The ratio between the difference of the first and second magnitude values and the sum (*I*_*DSR*_) satisfies these requirements:

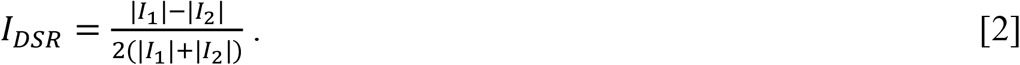

Multiplying the denominator by 2 ensures that *I*_*DSR*_-values fall within the same range as the *I*_*UNI*_-values. As *I*_*DSR*_ equals ±0.5 when either |*I*_*1*_| or |*I*_*2*_| is 0, and *I*_*DSR*_ equals 0 where |*I*_*1*_|=|*I*_*2*_|, the *I*_*DSR*_ contrast is indeed distinct from *I*_*UNI*_, where this behaviour is inverted. Similar to *I*_*UNI*_, nuisance factors contributing equally to |*I*_*1*_| and |*I*_*2*_| are cancelled out in *I*_*DSR*_.

Successful non-bijective mapping is performed by using a 2D-lookup table (2D-LUT), connecting a pair of (*I*_*UNI*_, *I*_*DSR*_)-values to their corresponding *R*_1_-value, henceforth referred to as 2D-LUT-MP2RAGE. In its simplest implementation, the 2D-LUT procedure compares the Euclidean distance between measured and reference value-pairs to obtain the *R*_1_-value (here pdist2 in MATLAB; The MathWorks, Inc., Natick, MA); this is illustrated in Figure 2. To ensure that the *R*_1_-estimation is driven by *I*_*UNI*_-values, as is the case for 1D-LUT, a larger weight can be applied to *I*_*UNI*_ by equally scaling the measured and reference *I*_*UNI*_-values.

**Figure 2:**
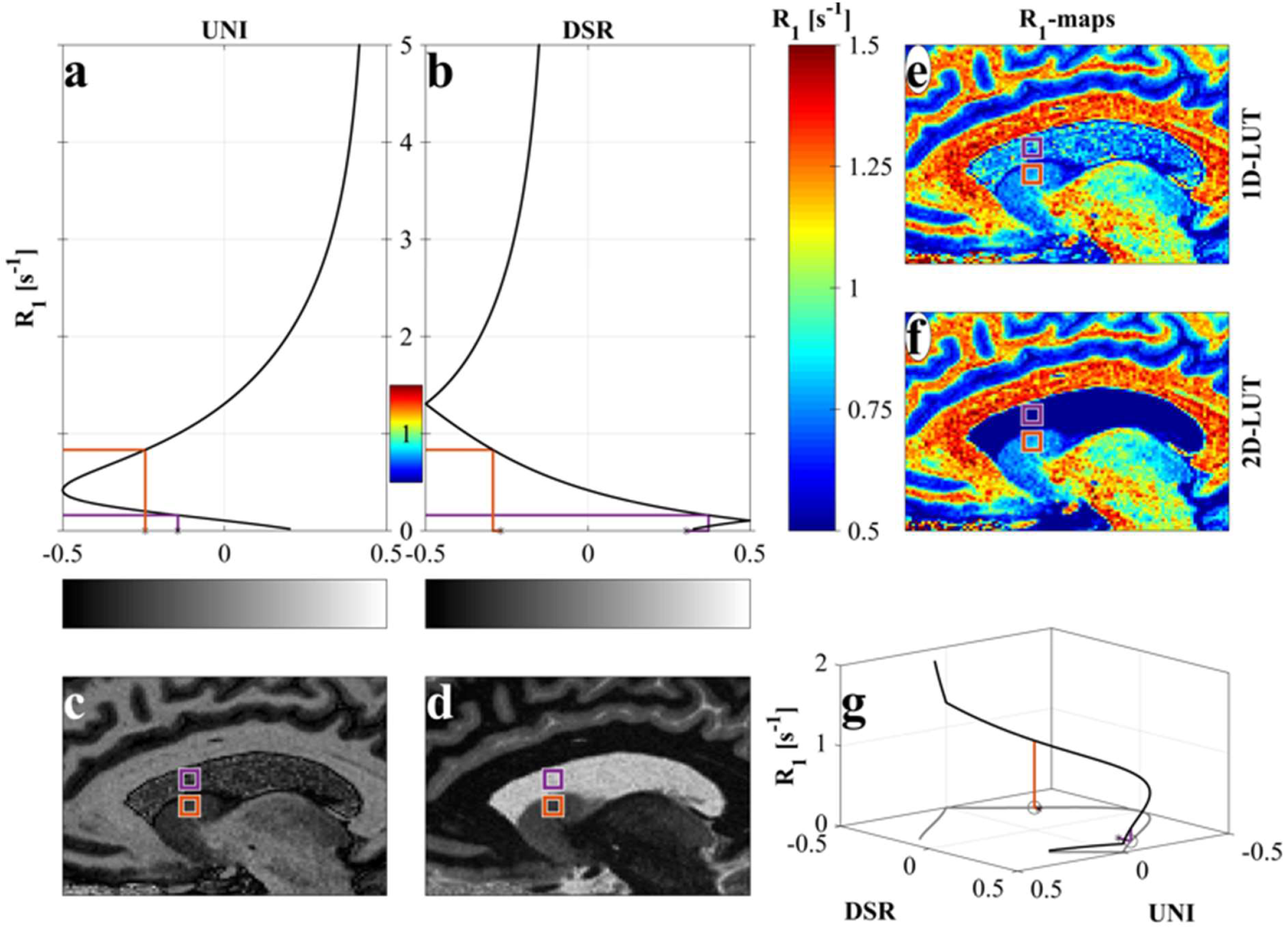
(a,b) 1D transfer curves for the MP2RAGE UNI (c) and DSR (d) of the LessBias protocol, respectively. (e,f) Resulting 1D- and 2D-LUT R_1_-maps, respectively. (g) 2D transfer curve between MP2RAGE UNI, DSR and R_1_. The MP2RAGE DSR, (d), can be used to disentangle the ambiguous relationship between the MP2RAGE UNI image, (c), and the R_1_-value, (a,b). The method is illustrated for CSF (purple) and deep GM (orange) voxels with MP2RAGE UNI values close to each other. By Identifying the MP2RAGE values in both the MP2RAGE UNI image and the MP2RAGE DSR image, a unique R_1_-value can be assigned to each voxel using a 2D-LUT (f), but not a 1D-LUT (e). (g) Illustrates that the mapping is unambiguous in the full MP2RAGE UNI, DSR, R_1_-space.

## 3. Methods

The purpose of the present study is twofold: first, to verify that *R*_1_-mapping can be performed successfully in expanded sequence parameter space, by resolving non-bijective mapping through 2D-LUT; and second, that from this expanded parameter space, parameters can be selected that result in *R*_1_-maps unaffected by *B*_1_^+^-inhomogeneities to an extent unseen without non-bijective mapping. To demonstrate our proposed method, we acquired experimental data in multiple sessions of all authors (Subject 1-3) on a 3T Siemens PrismaFit system (Siemens Healthineers, Erlangen, Germany), using a 32-channel receive-only head coil. Images were acquired with the default configuration of the host site (Standard), and a new protocol (LessBias) with improved tolerance to *B*_1_^+^-inhomogeneities in the final *R*_1_-maps: 1) Standard: *T*_R,MP2RAGE_=5s, *T*_R,FLASH_=7.18ms, FA_1_/FA_2_=4º/5º, *T*_I,1_/*T*_I,2_=700ms/2500ms; 2) LessBias: *T*_R,MP2RAGE_=5s, *T*_R,FLASH_=7.18ms, FA_1_/FA_2_=3º/5º, *T*_I,1_/*T*_I,2_=500ms/1900ms. In both cases, a single acquisition took 11:20 minutes with a 215.6×172.8×230mm field of view (Anterior-Posterior/Left-Right/Superior-Inferior; phase/slice/read), grid size 256×192×240, i.e. 0.9mm^3^ isotropic voxels, partial Fourier=6/8, and 2xGRAPPA^11^. FatNavs motion correction was applied^12,13^. To systematically modify *B*_1_^+^-variation, we modified the excitation pulse amplitude (“SRFExcit” in the scan card; Supporting Figure S1). Each protocol was acquired three times, with SRFExcit at 60%, 100% and 140% of the reference voltage, which reflect the range of *B*_1_^+^-inhomogeneities seen beyond 3T, e.g., at 7T^4^.

Following FatNavs motion correction^12,13^, the anterior and posterior commissure (AC/PC) were identified on the Standard 100%-excitation pulse amplitude image. Next, all other images were reoriented accordingly, to obtain a near AC-PC space orientation in all acquisitions. For each acquisition, an *R*_1_-map was created using 1D-LUT^10^ and 2D-LUT for Standard and LessBias acquisitions, respectively.

The 2D-LUT *R*_1_-mapping was performed by using a step-size of Δ*R*_1_=0.001s^-1^ from *R*_1_=0.01s^-1^ to 20s^-1^. These hyperparameters were chosen empirically to avoid 2D-LUT truncation effects. We opted for equidistant *R*_1_-spacing over equidistant *T*_1_-spacing to better capture variations in the WM-GM range. To compare 1D-LUT and 2D-LUT *R*_1_-mapping most directly, we applied a strong, empirically determined 100:1 weight ratio between *I*_*UNI*_ and *I*_*DSR*_ before calculating the Euclidian distance. Consequently, *R*_1_-values were assigned on the basis of the *I*_*UNI*_ values, with *I*_*DSR*_ values used primarily for disambiguation.

The resultant *R*_1_-maps were segmented using SPM12 r7711^14,15^, and from the segmented images, two masked *R*_1_-maps were created. The first masked image was constructed from a 1-step eroded brain mask containing voxels where the sum of posterior probabilities for CSF, GM and WM exceeded 0.2. The second was constructed similarly, but included skull and soft tissue voxels. The brain-only *R*_1_-maps were co-registered to the Standard 100%-excitation pulse amplitude acquisition, and the estimated registrations were applied to all images. From these brain-masked, co-registered *R*_1_-maps, mean and standard deviation maps across the different excitation pulse amplitude scalings for both protocols (Standard and LessBias) were calculated. These mean and standard deviation maps were used to assess the standard deviation within each tissue type and visualise the *B*_*1*_^*+*^-effects in WM and CSF.

The effects within GM were visualised on a central GM surface. Central GM surfaces were constructed using CAT12 r2560^16–19^ to segment the head-masked, co-registered *R*_1_-maps with the iso-volume approach^20,21^. By sampling each *R*_1_-map on their corresponding central GM surface and projecting out the local curvature^1^, we obtained curvature-corrected central GM *R*_1_-surfaces for each image. Prior to curvature correction, both curvature- and *R*_1_-surfaces were resampled to the FreeSurfer template surface in CAT12, and 2D-smoothed with a Gaussian kernel (FWHM 6mm). From these surfaces, we calculated the standard deviation across the different excitation pulse amplitude scalings, for both protocols. The curvature-corrected *R*_1_-values and their standard deviations were then visualised on an inflated GM template surface.

To compare the effects of noise and *B*_*1*_^*+*^-inhomogeneity on *R*_*1*_, we acquired three images for both protocols in one subject without SRFExcit manipulations. Instead, the nominal FA provided during *R*_1_-mapping was manipulated in steps of 10%. Ten *R*_1_-maps were derived for each image, where FA for one image was left untouched, FA for one image was reduced 0-90%, and the FA in the last image was increased 0-90%. Preprocessing of these images was left as described above.

The Supporting Information (overview in Supporting Table S1.) further describes quantitative *R*_1_-accuracy assessment with respect to curvature correction (section 4), segmentation robustness (section 5), mean *R*_1_-values across subjects, protocols and sessions (section 6), FatNavs effects (section 6), and influence of number of slices (section 5 and 6).

In addition, we acquired SA2RAGE images during each session to compare the efficacy of LessBias against post-processing *B*_*1*_^*+*^-map correction^22^. *B*_*1*_^*+*^-correction was applied to the Standard images using in-house code (see Supporting Information, section 7), and the corrected *R*_*1*_-maps were compared to both uncorrected-Standard and LessBias *R*_1_-maps (see Supporting Figures S11-S15).

## 4. Results

SNR between Standard and LessBias is highly comparable for nominal excitation FAs, i.e., SRFExcit of 100%. When SNR is defined as the maximum intensity value across |*I*_*1*_| and |*I*_*2*_| divided by the mean intensity value of the background of |*I*_*1*_|, SNR_Standard_≈75 and SNR_LessBias_≈71.

Figure 1 shows the transfer curves of Standard (1a) and LessBias (1b), the latter showing non-bijection within the *R*_1_-window-of-interest (0.2-5s^-1^), with the fold occurring at approximately *R*_1_=0.4s^-1^. Additionally, the estimated *B*_1_^+^-inhomogeneity tolerance in the GM and white matter (WM) range for LessBias is strongly improved compared to Standard. Figure 2 visualises the 2D-LUT-MP2RAGE. Figure 2e shows how the 1D-LUT approach gives rise to ambiguous *R*_1_-values across CSF (purple square) and GM (orange square). Figure 2f shows how 2D-LUT resolves the non-bijective mapping.

Figures 3 and 4 outline the results of amplifying *B*_1_^+^ across acquisitions. In Figure 3, substantial WM *R*_1_-variation is seen across the different levels of *B*_1_^+^-amplification in Standard for Subjects 1-3. In contrast, LessBias combined with 2D-LUT, provide consistent WM *R*_1_-values across *B*_1_^+^-amplifications. For a detailed overview of the standard deviations across protocols, see Supporting Tables S5-7.

**Figure 3:**
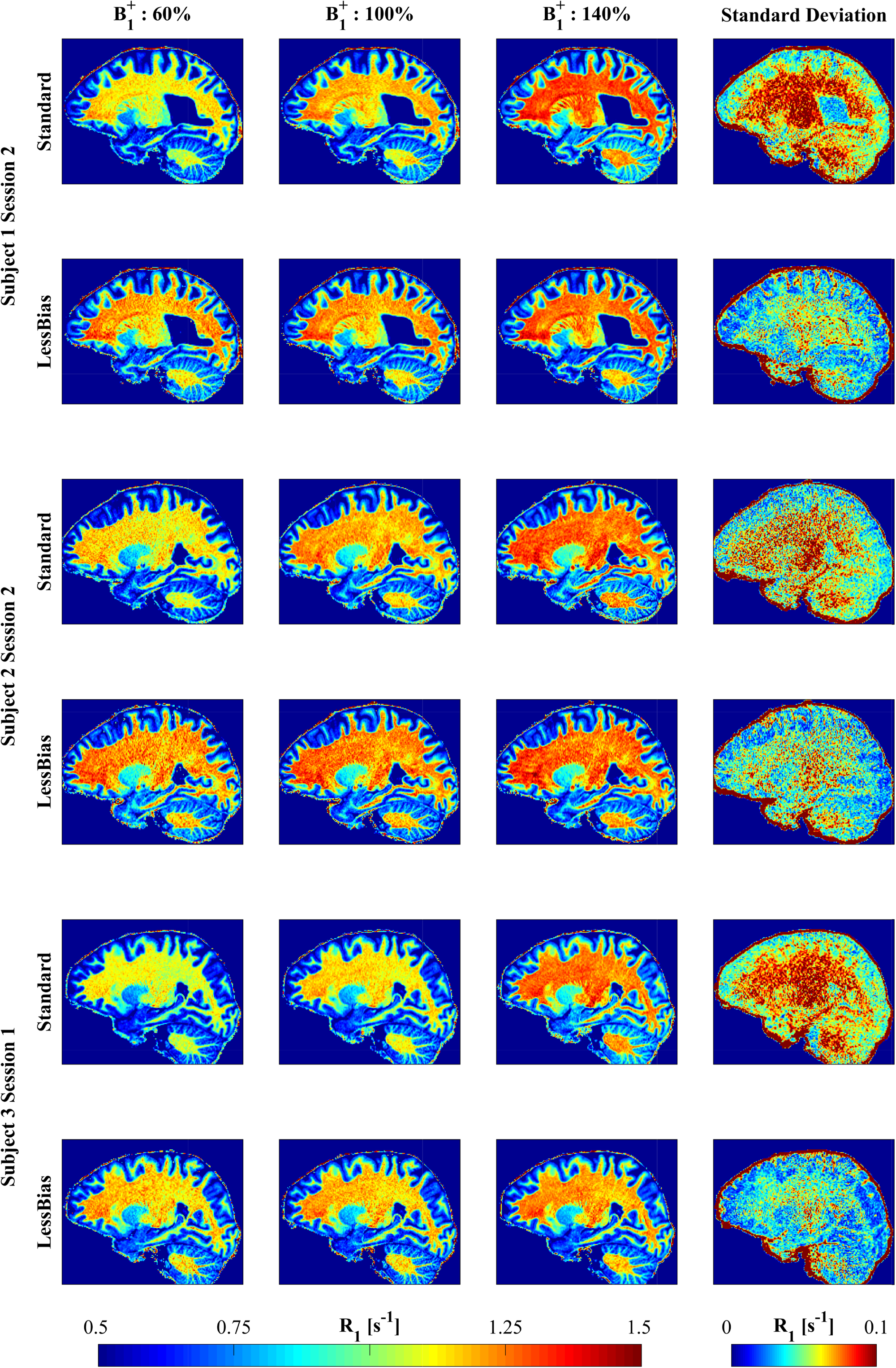
Volumetric R_1_-maps from the Standard (odd rows) and LessBias (even rows) protocol for Subject 1 (top), 2 (middle) and 3 (bottom). By scaling the excitation pulse amplitude to 60%, 100% and 140% (columns 1-3, respectively), it is possible to visualise the benefit of the LessBias protocol. As can be seen from both the individual acquisitions and the standard deviation image (column 4), the LessBias protocol does indeed, as predicted from Figure 1, demonstrate robustness to deviations from the nominal B ^+^. From the standard deviation images, it can be observed that LessBias provides a strongly reduced standard deviation for GM and WM, at the cost of a slightly increased standard deviation for CSF, compared to the Standard protocol. Actual mean and standard deviation of R_1_-values are listed in Supporting Tables S6.

In Figure 4, the variability in *R*_1_-values is displayed on central GM surfaces across Subjects 1-3. Standard (odd rows) shows substantial variation in *R*_1_-values compared LessBias (even rows). This is further illustrated by the standard deviation across each row (final column), where the standard deviations for LessBias are consistently substantially lower than for Standard. Highly myelinated areas (e.g. V1/V5-MT+/M1/S1/A1^1–3^) are reliably identifiable across LessBias images, where they become less distinct in Standard (e.g. V1 in Subject 1-Standard 140% or Subject 3-Standard 60%).

**Figure 4:**
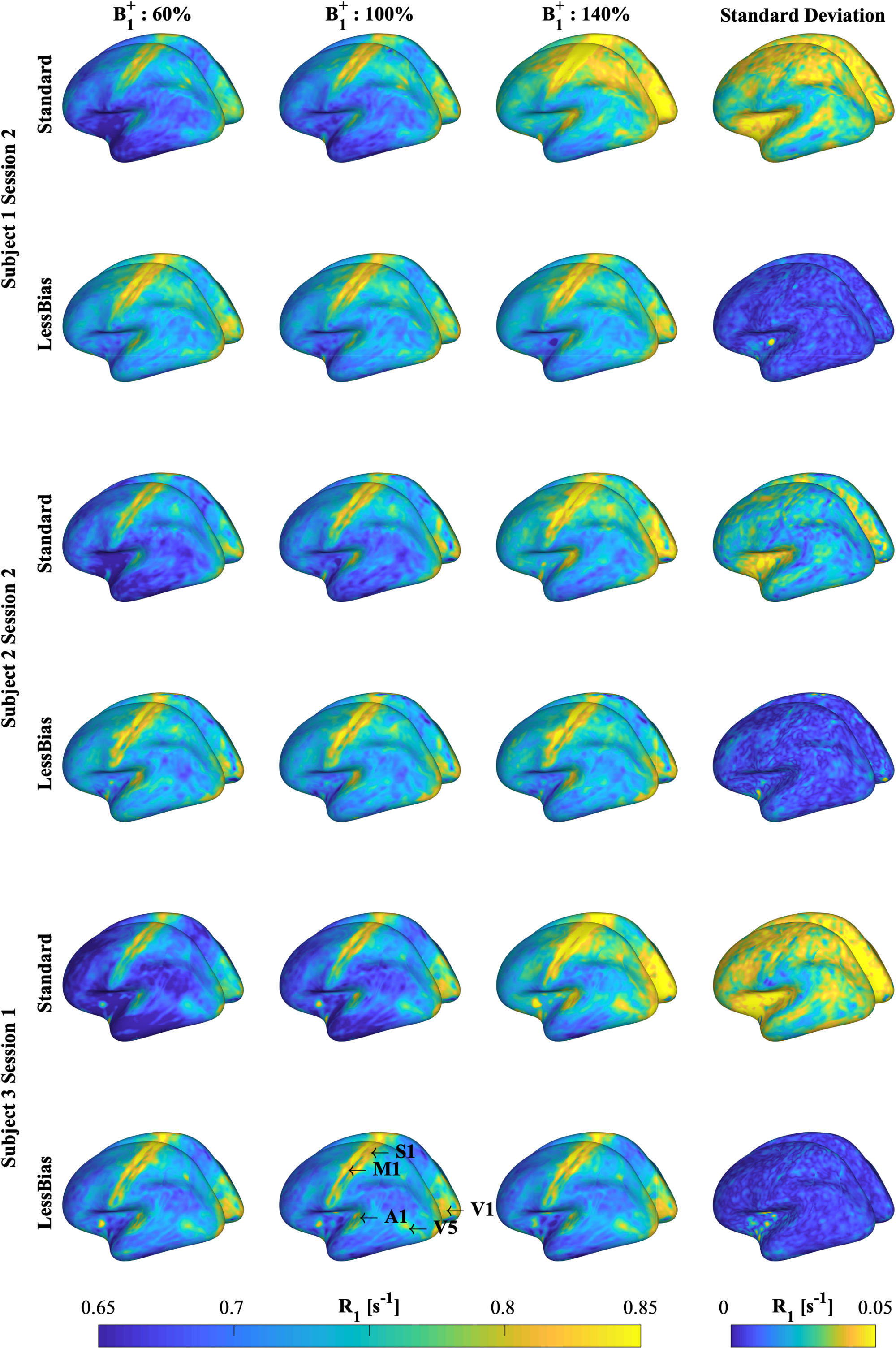
GM surface R_1_-maps from the Standard (odd rows) and LessBias (even rows) protocol for Subject 1 (top), 2 (middle) and 3 (bottom). GM surfaces are presented with uniform windowing across images (0.65 - 0.85 s^-1^). By scaling the excitation pulse amplitude to 60%, 100% and 140% (columns 1-3, respectively), it is possible to visualise the benefit of the LessBias protocol. As can be seen from both the individual acquisitions and the standard deviation surface (column 4), the LessBias protocol does indeed, as predicted from Figure 1, demonstrate robustness to deviations from the nominal B_1_ ^+^. From the standard deviation surfaces, it can be observed that LessBias provides a strongly reduced standard deviation for GM, compared to the Standard protocol. Increased R_1_-values are observed in expected regions (V1, M1, S1, A1, and possibly V5). Cortical GM volume and thickness estimations are listed in Supporting Tables S3. Actual mean and standard deviation of R_1_-values are listed in Supporting Tables S6.

Figure 5 shows the results of manipulating the nominal FA during *R*_1_-mapping across comparable images from one session. At an FA-manipulation of ±0%, we expect the variance across images to be driven primarily by noise. Differences in standard deviation at ±0% and the FA-manipulations indicate the contribution of bias.

**Figure 5:**
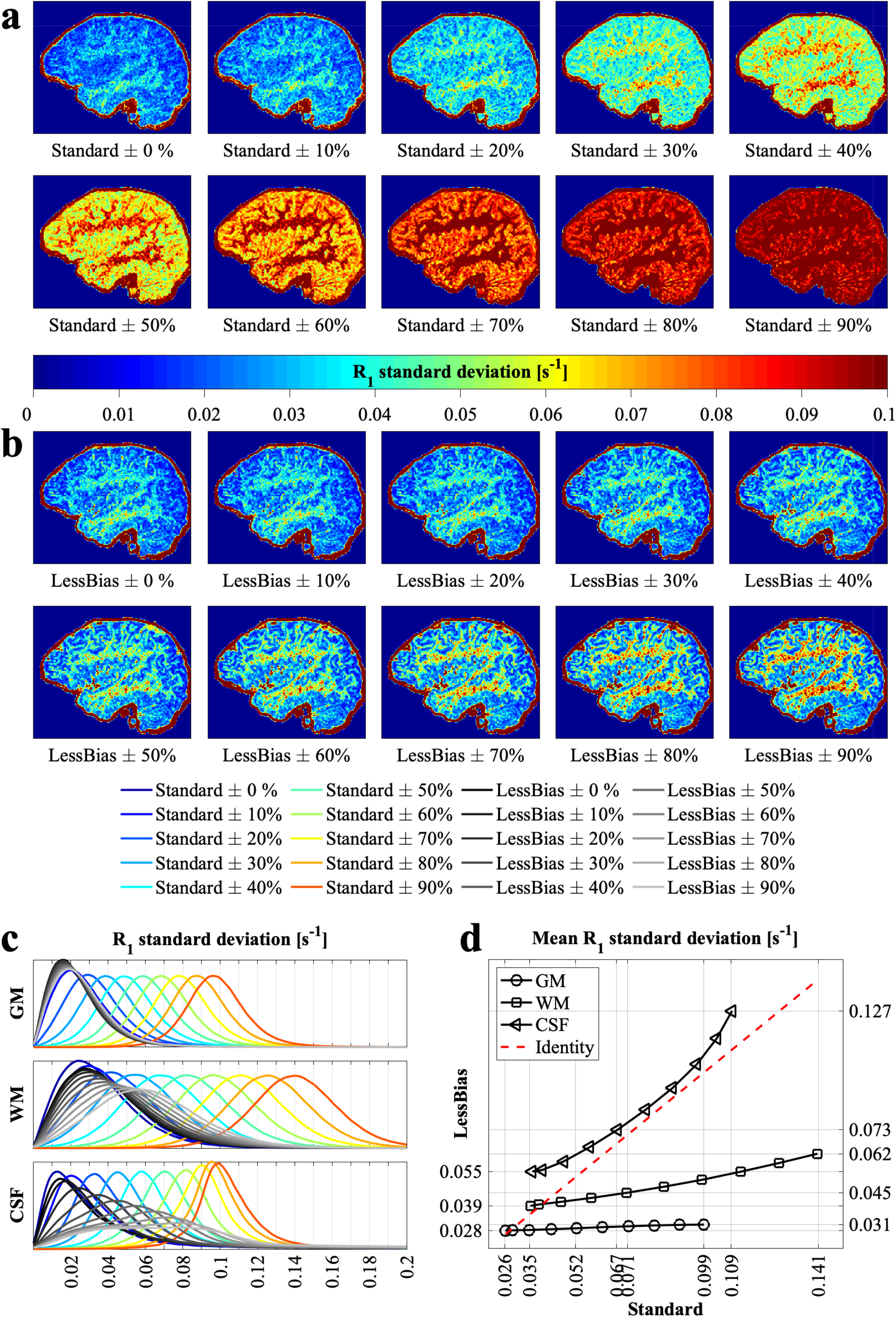
(a) R_1_ standard deviation maps of Standard for FA manipulations performed during R_1_-map estimation. (b) R_1_ standard deviation maps of LessBias for FA manipulations performed during R1-map estimation. (c) Distributions of voxel-wise R_1_ standard deviations per manipulation for Standard and LessBias (in colour and greyscale, respectively). For each tissue type (GM, WM and CSF), a mask was created from the intersections between Standard and LessBias of the segmented volumes from the unmanipulated (B_1_ ^+^: 100%) Standard and LessBias images. (d) Mean of the standard deviations distributions of (c) for GM, WM and CSF plotted against each other per manipulation. Leftmost/rightmost points belong to 0% and 90% manipulation, respectively. We expect that variation seen in (a) and (b) at ±0% is primarily driven by noise. Variation introduced as a result of the FA manipulation is assumed to be analogue to bias-induced variation. Visually, it appears in (a) that bias overtakes as the driving factor for variation at ±20%. This is confirmed by the histograms in (c), where especially in GM and WM, the distribution of standard deviations appears to be a linear shift based on the FA manipulation. This further becomes apparent, where we see that the contribution of an FA-manipulation of 20% and above greatly outweighs the contribution of a ±10% manipulation. Meanwhile, visual variation in LessBias for WM and especially GM remains minimal for a far higher FA-manipulation, as seen in (b). Distribution of standard deviations as a result of the manipulation in (c) are far less spread out in GM and WM for LessBias than Standard. In (d) we see that mean standard deviation in GM is increased by ∼10% at a ±90% FA-manipulation, and in WM ∼20% at a ±40% FA-manipulation (vs ∼300% and ∼75% in Standard respectively)

## 5. Discussion

With 2D-LUT-MP2RAGE *R*_1_-mapping, we have expanded the MP2RAGE sequence parameter space, enabling us to further refine the features of MP2RAGE acquisitions. In this paper, we focussed on the potential for improving *B*_1_^+^-inhomogeneity tolerance on the path towards ultrahigh resolution. Given the push for higher field strengths^23,24^, driven by this desire for higher resolutions, the problem of *B*_1_^+^-inhomogeneity intolerance will only become more pressing. To investigate levels of *B*_1_^+^-inhomogeneities similar to those seen at 7T, we manipulated the amplitude of the excitation pulse across images. It appears indeed possible to achieve greatly improved *B*_1_^+^-inhomogeneity tolerance with minimal SNR compromise. Using the adjusted sequence parameters in combination with 2D-LUT mapping, we drastically reduced the standard deviation of WM and GM *R*_1_-values without extending the acquisition time (see Supporting Tables S5-7).

Figure 5 showed that, for the Standard protocol, *B*_*1*_^*+*^-contributions to variance compared to noise is already substantial at ±20%. Meanwhile, Figure 5 suggests that LessBias is far less susceptible to *B*_*1*_-effects in GM and WM, showing nearly no effect of bias in GM and a drastically reduced effect in WM compared to Standard. Similarly, variability in CSF is greatly reduced in LessBias up to a point, showing a shift at around 40%. This shift could be because the analysis captures consistency, not accuracy. Beyond a certain FA-manipulation, variability may top out, slowing down the trajectory of Standard. It should also be noted that the SNR during each manipulation remains constant, while variations in the FA during acquisition would result in altered SNRs. As a result, Figure 5 cannot catch the total effect of *B*_*1*_^*+*^-inhomogeneities, but only the variation driven by deviations from the nominal transfer curve.

There are alternative methods that correct for *B*_1_^+^ either in post-processing^25,26^, (e.g., SA2RAGE^22^ or MP3RAGE^27^), or at the acquisition stage, e.g., pTx^5,6^ and Universal Pulses (UPs)^7–9^. For example, pTx PASTeUR UPs substantially reduced *B*_1_^+^-inhomogeneities at 7T compared to Circular Polarization mode^8^. The remaining *B*_1_^+^-inhomogeneity should be manageable with 2D-LUT-MP2RAGE. Note that the parameters used in the protocol can be adjusted to the user’s needs, but at the expense of other parameters. For instance, number of slices can be increased for higher resolution imaging, but to maintain *B*_1_^+^-inhomogeneity tolerance, requires additional adjusting of *T* ’s and FAs. The extent of these adjustments would further depend on what other *B*_1_^+^-inhomogeneity compensation techniques are available and the *B*_*0*_. For an example, see Supporting Information section 8.

Whether or not MP2RAGE is acquired in combination with e.g. UPs, we see no immediate downsides of utilising 2D-LUT-MP2RAGE, given that the potential benefits come at virtually no cost compared to standard MP2RAGE applications.

On its own, 2D-LUT-MP2RAGE provides *B*_1_^+^-inhomogeneity tolerance at the voxel-level without modified sequences or increased scan times, and with sufficient time for fat navigators. Time otherwise needed for such high-resolution *B*_1_^+^-inhomogeneity corrections can instead be utilised to improve throughput, SNR or resolution of *R*_1_-images. Given that 2D-LUT-MP2RAGE is agnostic to hardware setup and availability of secondary sequences, we anticipate that it can be implemented to tackle the challenges of cross-site collaborations^28^, to a larger extent than alternative methods.

Recent developments with synthetic *T*_1_-contrasts showed that through a sufficiently accurate *T*_1_-map, different *T*_1_-weighted contrasts can be derived in post-processing, and that this can be used to optimise scan times^7^. When relying on synthetic MRI, the contrast of the original acquisition can be arbitrary, as long as the *T*_1_-mapping succeeds. This means that *I*_*UNI*_ no longer needs to be visually interpretable and permits non-bijection. Consequently, further optimisations for improved *B*_1_^+^-inhomogeneity tolerance, SNR, and/or acquisition times may be possible in the wider sequence parameter space. Desired (visually interpretable) contrasts can then be computed from the resultant *T*_1_-maps.

Other implementations for non-bijective transfer curves may be for specialised uses (e.g. Ref.^29^). Such cases may require sequence parameters that come with increased CNR in or across specific tissues (e.g. simultaneous brain and cervical spinal cord imaging). Reducing the slope of the transfer curve will lead to increased contrast between tissues in *I*_*UNI*_, but will also lead to non-bijective transfer curves when the slope drops below a certain threshold. When such protocols are desired, 2D-LUT-MP2RAGE will still be able to derive accurate *R*_1_-maps from the data. Through additional post-processing steps, the high contrast from the original images could be extracted, while the *R*_1_-map serves as a reference.

The remaining CSF *T*_1_-variance compromise impose limits e.g. in specific CSF-based MRI biomarkers and neurodegenerative-disease studies^30^. However, we stress that Standard suffered from stronger variance than LessBias. Through 2D-LUT-MP2RAGE, it may be possible to select parameters that address *T*_1_-variance in CSF, but this was not investigated. Additionally, Standard suffers from non-bijections around CSF at a low *B*_*1*_^*+*^, leading to larger misestimations in 1D-LUT that cannot be corrected with a 1D-LUT *B*_1_^+^-correction method.

This study assumed 96% inversion efficiency^4^, which anticipatedly is fairly valid at 3T. At ultrahigh field strength this assumption is likely not valid and investigations with respect to inversion efficiency and our method remain. This is where 2D-LUT-MP2RAGE may best be combined with UPs.

Our *B*_1_^+^-maps (e.g. Supporting Figure S16) show an expected ±20% variation, but we stress-tested our method with ±40% excitation pulse amplitude scalings to mimic variations seen commonly at 7T^4^ as a feasibility test for future studies. This study does not directly answer how more severe variations are attacked, but as mentioned above we expect the method to add value in concert with other methods e.g. using Universal Pulses^8,9^.

## 6. Conclusions

With the 2D-LUT-MP2RAGE, we were able to open up the usable MP2RAGE acquisition parameter space, which can impart non-bijective MP2RAGE-to-*T*_1_ transfer curves. This increased freedom was used to address *B*_1_^+^-inhomogeneity without additional hardware or data than what the current MP2RAGE sequence inherently provides. Compared to typical MP2RAGE acquisition parameters, we saw greatly reduced variability across GM and WM *R*_1_-values with *B*_1_ ^+^-variations (±40%) mimicking those seen at 7T. While this paper focussed on improving *B*_1_^+^-inhomogeneity tolerance, this 2D-LUT mapping may also enable the use of protocols that provide contrast or scan times previously unseen in MP2RAGE.

## Code availability

Code and example dataset are available at https://github.com/torbenelund/2D-lookup-tools-for-MP2RAGE.

