## Supporting Information for "A 3T investigation of *B*_1_^+^-inhomogeneity tolerance in MP2RAGE-based *R*_1_-mapping by calculating second contrast that permits previously problematic sequence parameters"

September 6, 2024

#### Contents


List of Figures 1

List of Tables 5

1 Overview of acquisitions 8

2 How to change excitation flipangle to modify $B_1^+$ 9

3 $R_1$-nulling 10

4 The impact of curvature correction on $R_1$-maps 11

5 Robustness of CAT12 segmentation outputs 13

6 Robustness of $R_1$-values 16

7 Effect of $B_1^+$-correction. 26

8 A 7T example with 512 slices 34


#### List of Figures


S1 Modifying the excitation FA. Screenshot of Syngo scan card user interface (Siemens Healthineers, Erlangen, Germany) showing in orange highlights where SRFExcit was modified. . . . . 9

S2 $R_1$-nulling. Distribution of voxel-wise standard deviations across excitation pulse scalings over the mean $R_1$for the Standard (top) and LessBias (bottom) protocols. Increased density of the distribution is seen for $R_1 \approx 0.7\text{s}^{-1}$ (GM) and $R_1 \approx 1.15\text{s}^{-1}$ (WM) at lower standard deviations in LessBias compared to Standard, suggesting higher consistency across the $R_1$-estimations. Voids (white) in both distributions correspond with folds in the respective transfer curves ($R_1 < 0.26\text{s}^{-1}$ in Standard; $R_1 \approx 0.1\text{s}^{-1}$ (fold in the DSR transfer curve) or $R_1 \approx 0.4\text{s}^{-1}$ (fold in the UNI transfer curve) in LessBias), indicating $R_1$-nulling. 10

S3 The effect of curvature correction on the estimated $R_1$-surfaces and the standard deviation across $B_1^+$manipulations. By projecting out the local curvature of the $R_1$-values it is possible to remove curvature-induced variation in the $R_1$-mapping. As seen from the two bottom rows, the estimated curvatures are almost identical across protocols and $B_1^+$-manipulations. From the standard deviation of the curvature-uncorrected $R_1$-maps (two top rows) and curvature-corrected $R_1$-maps (two central rows), it is seen that these are entirely driven by the $B_1^+$-manipulations. . . . . . . . . . . . . . . . . . . . . . . . . . . . . 12

S4 GM $R_1$-assessment, Subject 1 Session 3, regarding multiple repetitions. Panel a) shows distributions of $R_1$-values on the central GM-surface for both Standard and LessBias, across three repetitions within the same scanning session. In panel b) the surfaces corresponding to the curves in panel a) are shown. In panel c) 2D histograms show the relationship between Standard and LessBias central, cortical GM $R_1$-values. The red lines indicate the perfect relationship between the data and the dashed, black lines indicate the best linear fit with an intercept of zero. . . . . . . . . . . . . . . . . . . . . . . . . . . . . . . . . . . . . . . 20

S5 GM $R_1$-assessment, Subject 1 Session 1-3, regarding multiple sessions. Panel a) shows distributions of $R_1$-values on the central GM-surface for both Standard and LessBias, across three scanning sessions. In panel b) the surfaces corresponding to the curves in panel a) are shown. In panel c) 2D histograms show the relationship between the Standard and the LessBias central, cortical GM $R_1$-values. The red lines indicate the perfect relationship between the data and the dashed, black lines indicate the best linear fit with an intercept of zero. . . 21

S6 GM $R_1$-assessment, Subject 2 Session 1-3, regarding multiple sessions. Panel a) shows the distribution of $R_1$-values on the central GM-surface for both Standard and LessBias across three scanning sessions. Panel b) shows the surfaces corresponding to the curves in panel a). Panel c) shows 2D histograms of the relationship between Standard and LessBias central GM cortical $R_1$-values. It is seen that both Standard and LessBias are quite consistent across sessions, although not quite at the level seen in subject1 Figure S5 where a less motion was present. See Table S5 for a whole brain comparison. . . . . . . . . . . . . 22

S7 GM $R_1$-assessment, Subject 3 Session 1, regarding multiple $B_1^+$-values. Panel a) shows distributions of $R_1$-values on the central GM-surface for both Standard and LessBias, across three $B_1^+$-manipulations. From the distributions and their median values (indicated with vertical lines) it is clear that LessBias method is quite consistent across $B_1^+$-values, but Standard fluctuates rapidly. Panel b) shows the surfaces corresponding to the curves in panel a). The Standard also visually shows a substantial fluctuation in $R_1$-values, whereas LessBias shows much less variation. Panel c) shows 2D histograms of the relationship between Standard and LessBias central GM cortical $R_1$-values. It is overall seen that LessBias presents with more constant distributions across the $B_1^+$-manipulations than Standard, and that Standard agrees most with LessBias when $B_1^+$: 100 %. 23
S8 GM $R_1$-assessment, Subject 2 Session 1, regarding number of slices. Panel a) shows distributions of $R_1$-values on the central GM-surface for both Standard and LessBias for the different number of slices. Panel b) shows the surfaces corresponding to the curves in panel a). Panel c) shows 2D histograms of the relationship between Standard and LessBias central GM cortical $R_1$-values. It is seen that in the examined interval, both protocols seem quite robust to changes in the number os slices. See Table S7 for a whole brain comparison. . . . . . . . 24
S9 GM $R_1$-assessment, Subject 2 Session 3, regarding the fat navigator module. Panel a) shows distributions of $R_1$-values on the central GM-surface for both Standard and LessBias, with and without a fat navigator module present in the sequence. To be clear, "FatNavs" indicates that the acquisition was performed using the FatNavs-MP2RAGE sequence (Gallichan et al. MRM 75 (2016) 1030-1039), whereas "NoFatNavs" indicates that the MP2RAGE product version without fat navigators was used. The data in this analysis from the "FatNavs" scans, however, was the non-motion corrected data set. From the distributions (median values indicated with vertical lines) it is clear that the fat navigator pulse have no effect by itself. Panel b) shows the surfaces corresponding to the curves in panel a). In panel c), 2D histograms also show the lack of effect from the fat navigator module. The red lines indicate the perfect relationship between the data and the dashed, black lines indicate the best linear fit with an intercept of zero. An almost perfect correlation between the two MP2RAGE protocols is seen for both LessBias and Standard, indicating that the FatNavs pulse did not, by it self, influence the estimated $R_1$-values. . . . . . . . . . . . . . . . . . . . . . 25
S10 Phantom $B_1^+$-maps. Instead of the original, ratio-based SA2RAGE method (three, top left images), we estimated our $B_1^+$-maps (three, top right images) through the $I_{\text{UNI}}$-values and performed the correct via a 3D-LUT procedure. The essential difference between the two procedures is shown in the three bottom left images. . . . . . . . . . . . . . . . . . . . . . . . . . . . . . . . . . . . 27

S11 GM $R_1$-assessment, Subject 1 Session 2, regarding $B_1^+$-correction. Panel a) shows the distribution of $R_1$-values on the central GM-surface for both Standard and LessBias, As well as for a $B_1^+$-corrected version of Standard. From the distributions and their median values (indicated with vertical lines) it is clear that the $R_1$-values from Standard resemble those of LessBias, when the $R_1$-map from Standard is $B_1^+$-corrected. Panel b) shows the surfaces corresponding to the curves in panel a). A shift towards higher $R_1$-values at the cortex is seen, when the $R_1$-map from Standard is $B_1^+$-corrected. Panel c) shows 2D histograms for the three possible combinations of the maps in panel b). The red lines indicate the perfect relationship between the data and the dashed, black lines indicate the best linear fit with an intercept of zero. It is seen from the plots that while the maps from LessBias and Standard are fairly similar, the best correspondence is found between the $R_1$-maps from $B_1^+$-corrected Standard and LessBias. . . . . 28
S12 GM $R_1$-assessment, Subject 1 Session 3, regarding $B_1^+$-correction. Panel a) shows the distribution of $R_1$-values on the central GM-surface for both Standard and LessBias, As well as for a $B_1^+$-corrected version of Standard. From the distributions and their median values (indicated with vertical lines) it is clear that the $R_1$-values from Standard resemble those of LessBias, when the $R_1$-map from Standard is $B_1^+$-corrected. Panel b) shows the surfaces corresponding to the curves in panel a). A shift towards higher $R_1$-values at the cortex is seen, when the $R_1$-map from Standard is $B_1^+$-corrected. Panel c) shows 2D histograms for the three possible combinations of the maps in panel b). The red lines indicate the perfect relationship between the data and the dashed, black lines indicate the best linear fit with an intercept of zero. It is seen from the plots that while the maps from LessBias and Standard are fairly similar, the best correspondence is found between the $R_1$-maps from $B_1^+$-corrected Standard and LessBias. . . . . 29
S13 GM $R_1$-assessment, Subject 2 Session 1, regarding $B_1^+$-correction. Panel a) shows the distribution of $R_1$-values on the central GM-surface for both Standard and LessBias, As well as for a $B_1^+$-corrected version of Standard. From the distributions and their median values (indicated with vertical lines) it is clear that the $R_1$-values from Standard resemble those of LessBias, when the $R_1$-map from Standard is $B_1^+$-corrected. Panel b) shows the surfaces corresponding to the curves in panel a). A shift towards higher $R_1$-values at the cortex is seen, when the $R_1$-map from Standard is $B_1^+$-corrected. Panel c) shows 2D histograms for the three possible combinations of the maps in panel b). The red lines indicate the perfect relationship between the data and the dashed, black lines indicate the best linear fit with an intercept of zero. It is seen from the plots that while the maps from LessBias and Standard are fairly similar, the best correspondence is found between the $R_1$-maps from $B_1^+$-corrected Standard and LessBias. . . . . 30

S14 GM $R_1$-assessment, Subject 2 Session 2, regarding $B_1^+$-correction. Panel a) shows the distribution of $R_1$-values on the central GM-surface for both Standard and LessBias, As well as for a $B_1^+$-corrected version of Standard. From the distributions and their median values (indicated with vertical lines) it is clear that the $R_1$-values from Standard resemble those of LessBias, when the $R_1$-map from Standard is $B_1^+$-corrected. Panel b) shows the surfaces corresponding to the curves in panel a). A shift towards higher $R_1$-values at the cortex is seen, when the $R_1$-map from Standard is $B_1^+$-corrected. Panel c) shows 2D histograms for the three possible combinations of the maps in panel b). The red lines indicate the perfect relationship between the data and the dashed, black lines indicate the best linear fit with an intercept of zero. It is seen from the plots that while the maps from LessBias and Standard are fairly similar, the best correspondence is found between the $R_1$-maps from $B_1^+$-corrected Standard and LessBias. . . . . 31

S15 GM $R_1$-assessment, Subject 3 Session 1, regarding $B_1^+$-correction. Panel a) shows the distribution of $R_1$-values on the central GM-surface for both Standard and LessBias, As well as for a $B_1^+$-corrected version of Standard. From the distributions and their median values (indicated with vertical lines) it is clear that the $R_1$-values from Standard resemble those of LessBias, when the $R_1$-map from Standard is $B_1^+$-corrected. Panel b) shows the surfaces corresponding to the curves in panel a). A shift towards higher $R_1$-values at the cortex is seen, when the $R_1$-map from Standard is $B_1^+$-corrected. Panel c) shows 2D histograms for the three possible combinations of the maps in panel b). The red lines indicate the perfect relationship between the data and the dashed, black lines indicate the best linear fit with an intercept of zero. It is seen from the plots that while the maps from LessBias and Standard are fairly similar, the best correspondence is found between the $R_1$-maps from $B_1^+$-corrected Standard and LessBias. . . . . 32

S16 GM $R_1$-assessment, Subject 2 Session 1, regarding $B_1^+$-correction. Panel a) shows distributions of $R_1$-values on the central GM-surface for both protocols, with and without $B_1^+$-correction. From the distributions and their median values (indicated with vertical lines) it is clear that the $R_1$-values from Standard resemble those of LessBias, when the $R_1$-map from Standard is $B_1^+$-corrected. However, as could be expected from Figure 5, the effect of $B_1^+$-correction on LessBias is minimal. Panel b) shows the surfaces corresponding to the curves in panel a). A shift towards higher $R_1$-values at the cortex is seen, when the $R_1$-map from Standard is $B_1^+$-corrected. In panel c), 2D histograms show the effect of $B_1^+$-correction for the two different protocols. The red lines indicate the perfect relationship between the data and the dashed, black lines indicate the best linear fit with an intercept of zero. It is seen that LessBias and LessBias with $B_1^+$-correction have a stronger correlation than Standard and Standard with $B_1^+$-correction. Panel d) shows the $B_1^+$-map used for the correction. . . . . . . . . . . . . . . . . . . . . 33

S17 Transfer functions for two protocols **a**: Standard* and **b**: LessBias* modified to acquire 512 slices at 7T. . . . . . . . . . . . . . . . . . . . . . . . . . . . . . 34

### List of Tables

S1 Supporting Figures overview with respect to investigated features. A • indicates a feature that has been investigated. . . . . . . . . . . . . . . . . . . . . . . . . 8

S2 Segmentation robustness within and across sessions. Top (Subject 1 Session 3) shows the variation of estimated volumes across repeated acquisitions of the two protocols. It is seen that LessBias shows a 2-3 fold larger standard deviation in estimated volumes, when compared to Standard, but in both cases the standard deviation across repetitions are for GM approximately 0.25 % of the corresponding volumes. Center (Subject 1) and bottom (Subject 2) show that Standard yields a 2.68 and 1.7 times higher standard deviation in the GM volume estimation than LessBias, respectively, when compared different sessions of the same subject. . . . . . 13
S3 Segmentation robustness across subjects. GM, WM, CSF and TIV volumes and GM average thickness for different protocols, subjects and $B_1^+$-manipulations from four sessions in three subjects. It is consistently seen that while Standard shows substantial variation in estimated volumes across $B_1^+$-modifications, LessBias is practically immune to the $B_1^+$-alterations. The standard deviation seen in the estimated volumes for LessBias is at the level of that seen in the top of Table S4 where each protocol was repeated three times without $B_1^+$-modifications. For Standard on the other hand the standard deviation across $B_1^+$-alterations is up to 22 times larger than what is seen for LessBias. . . . . . 14
S4 Segmentation robustness across sequence parameters. Top (Subject 2 Session 1), it is seen that the estimated volumes for the two protocols change a lot when Standard is acquired with various number of slices, while this is not the case for LessBias. The standard deviation across slice modulations for Standard is 2-16 times greater than for LessBias, where the standard deviation is comparable to when no parameters are modified. In GM, the standard deviation of estimated GM volumes for Standard is almost 2% of the estimated volume. At the bottom (Subject 2 Session 3), the effect of $B_1^+$correction and presence of fat navigators (FatNavs) are shown. A $\bullet$ in the FatNavs column indicates that the acquisition was performed using the FatNavs-MP2RAGE sequence (Gallichan et al. MRM 75 (2016) 1030-1039), whereas a $\circ$ indicates that the product version without fat navigators was used. Both protocols were reconstructed on the scanner i.e. without performing the motion correction. For LessBias, the estimated volumes are practically unaffected by the fat navigators, while the standard deviation for Standard is 8 times larger than for LessBias. In GM, the standard deviation of estimated GM volumes for Standard is around 1% of the estimated volume. $B_1^+$-correction of Standard reduce the standard deviation between the protocols by a factor of 2. . . . . . 15
S5 $R_1$-values (mean and standard deviation) within and across sessions. . . . . . 17
S6 $R_1$-values (mean and standard deviation) across subjects. In all four sessions Standard and LessBias were subject to $B_1^+$manipulations through changes of the FA (60, 100, and 140 %), see Sec. 2. We see a substantially smaller $R_1$standard deviation for LessBias in GM and WM than for Standard. Improvements range from a factor of 3.3 to 21.5. For CSF, LessBias ranges from 2 to 1.04 times higher than that of Standard. . . . . . 18

S7 $R_1$-values (mean and standard deviation) across sequence parameters. In the top (Subject 2 Session 1), the number of slices were changed. In the bottom (Subject 2 Session 3) a • in the FatNavs column indicates that the acquisition was performed using the FatNavs-MP2RAGE sequence (Gallichan et al. MRM 75 (2016) 1030-1039), whereas a ◦ indicates that the product version without fat navigators was used. Both sequences were reconstructed on the scanner i.e. without performing the motion correction. All scans were obtained with the same 100 % FA setting, and we see that the standard deviations for all tissue types are below 0.007 $s^{-1}$ for both Standard and LessBias. . . . . . . . . . . . . 19

### 1 Overview of acquisitions

In order to explore the robustness of the 2D-LUT and the LessBias protocols a total of seven scanning sessions were performed on three subjects. In addition to the protocol presented in the paper, the following four variations were tested: 1) The possible impact of the curvature correction performed when sampling $R_1$-values at the central GM surface. 2) Three repeated acquisitions each of the two protocols (Standard and LessBias) without modifying the $B_1^+$at the stage of acquisition. This session (Subject 1, Session 3) was used to evaluate the relation between image noise and $B_1^+$-introduced bias. 3) Three different number of slices 160, 192 and 240 were used with each of the two protocols (Standard and LessBias). 4) Potential magnetisation transfer effect of fat navigators (FatNavs) were evaluated in a third session where both the Standard and LessBias protocol were acquired with and without fat navigators. An overview of the different sessions is presented in Table S1 below.

**Table S1:** Supporting Figures overview with respect to investigated features. A • indicates a feature that has been investigated.

| | Subject | Session | $B_1^+$-manip. | Repeat | #Slices | FatNavs | $B_1^+$-corr. | Curv.-corr |
|---|---|---|---|---|---|---|---|---|
| Figure S3 | 1 | 1 | ○ | ○ | ○ | ○ | ○ | • |
| Figure S4 | 1 | 3 | ○ | • | ○ | ○ | ○ | ○ |
| Figure S5 | 1 | 1,2,3 | ○ | ○ | ○ | ○ | ○ | ○ |
| Figure S6 | 2 | 1,2,3 | ○ | ○ | ○ | ○ | ○ | ○ |
| Figure S7 | 3 | 1 | • | ○ | ○ | ○ | ○ | ○ |
| Figure S8 | 2 | 1 | ○ | ○ | • | ○ | ○ | ○ |
| Figure S9 | 2 | 3 | ○ | ○ | ○ | • | ○ | ○ |
| Figure S11 | 1 | 2 | ○ | ○ | ○ | ○ | • | ○ |
| Figure S12 | 1 | 3 | ○ | ○ | ○ | ○ | • | ○ |
| Figure S13 | 2 | 1 | ○ | ○ | ○ | ○ | • | ○ |
| Figure S14 | 2 | 2 | ○ | ○ | ○ | ○ | • | ○ |
| Figure S15 | 3 | 1 | ○ | ○ | ○ | ○ | • | ○ |
| Figure S16 | 2 | 1 | ○ | ○ | ○ | ○ | • • | ○ |

#### 2 How to change excitation flipangle to modify $B_1^+$

If the scanner interface had allowed for non-integer flipangles (FAs) the ±40% modifications of $B_1^+$could have been implemented by just entering the corresponding FAs. As this is not possible, we modified the SRFExcit pulse amplitude as shown in Figure S1.

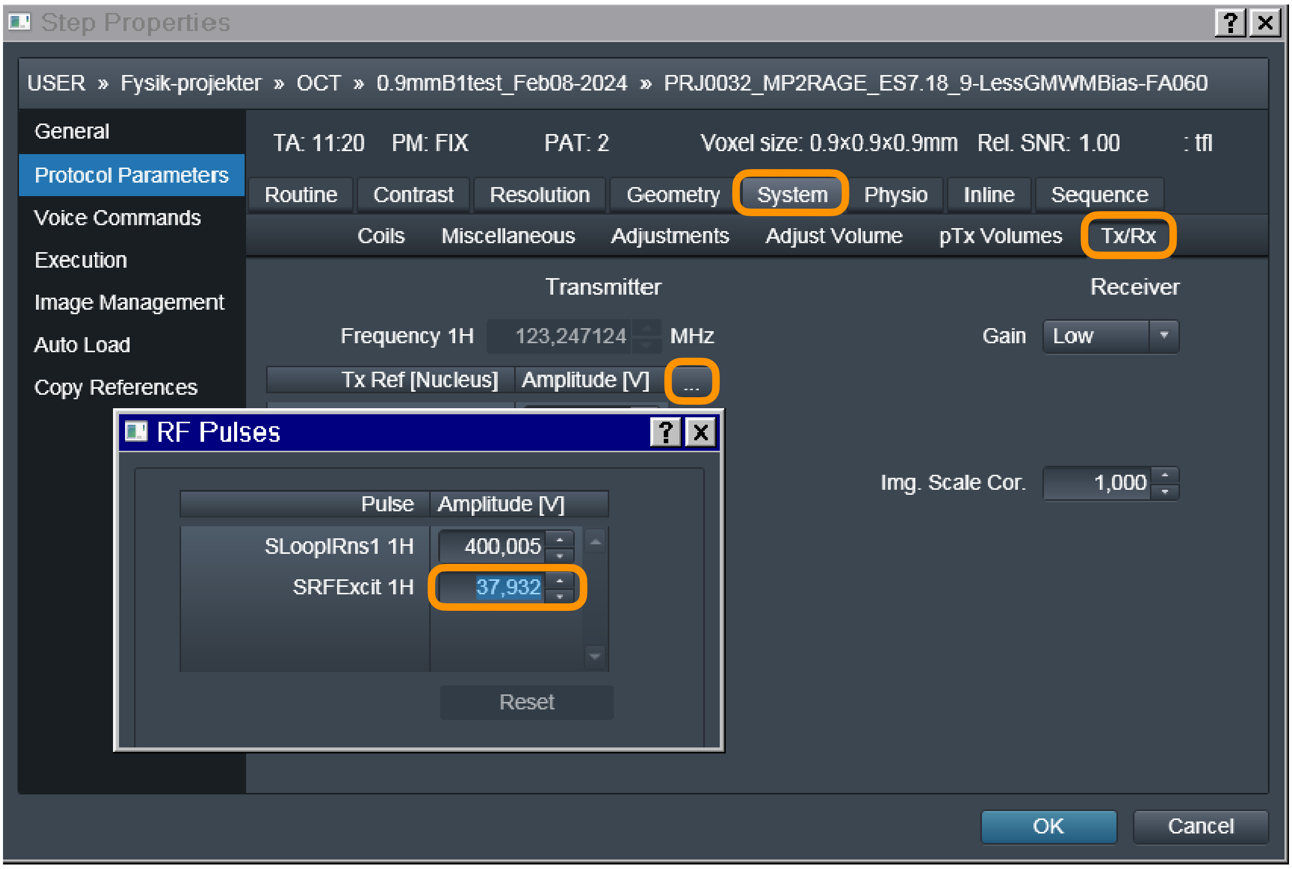


**Figure S1:** Modifying the excitation FA. Screenshot of Syngo scan card user interface (Siemens Healthineers, Erlangen, Germany) showing in orange highlights where SRFExcit was modified.

### 3 $R_1$-nulling

We observed a phenomenon we dubbed $R_1$-nulling, that is present in both 1D- and 2D-LUT $R_1$-mapping procedures. Certain $R_1$-values will not be assigned to voxels with certain characteristics. This can be seen in Figure S2, where white gaps appear in the distribution of $R_1$-values. Firstly, we noticed that this occurs for $R_1$-values close to the fold of the transfer curve (i.e. when $I_{\text{UNI}}$= -0.5). We speculate that this is the result of noise in the first and second inversion images. When computing the UNI values from these images, noise in either direction in the inversion images will necessarily always push the UNI-value closer to 0, whereas the UNI-values for voxels farther away from the boundaries of UNI can go either way as a result of noise. As a result, we expect that the SNR will have a substantial influence on the severity of this R1-nulling phenomenon. Both protocols avoid imposing this problem on the R1-maps by making sure $I_{\text{UNI}}$= -0.5 for $R_1$-values that are removed from tissue if interest (for Standard, at $R_1 \approx 0.12\text{s}^{-1}$, below CSF; for LessBias at $R_1 \approx 0.4\text{s}^{-1}$, between the expected $R_1$-values for CSF and GM). Note that for certain future purposes, like selective contrast, it may be difficult to avoid $R_1$-nulling in critical $R_1$-values, as a result of the location of the fold in the transfer curve or limitations of the SNR. More sophisticated algorithms may prove equipped to handle such data, and are under investigation.

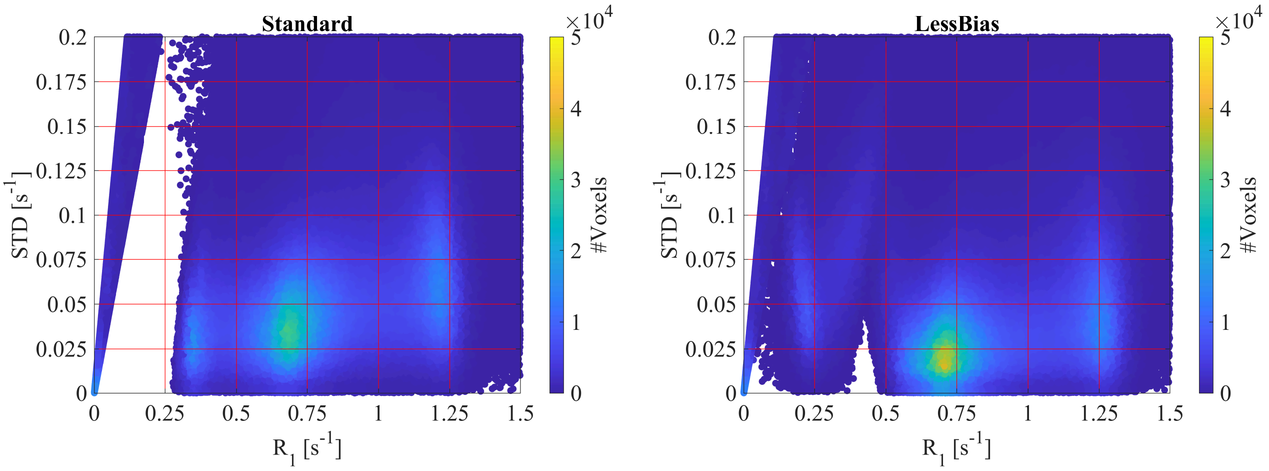


**Figure S2:** $R_1$-nulling. Distribution of voxel-wise standard deviations across excitation pulse scalings over the mean $R_1$for the Standard (top) and LessBias (bottom) protocols. Increased density of the distribution is seen for $R_1 \approx 0.7\text{s}^{-1}$ (GM) and $R_1 \approx 1.15\text{s}^{-1}$ (WM) at lower standard deviations in LessBias compared to Standard, suggesting higher consistency across the $R_1$-estimations. Voids (white) in both distributions correspond with folds in the respective transfer curves ($R_1 < 0.26\text{s}^{-1}$ in Standard; $R_1 \approx 0.1\text{s}^{-1}$ (fold in the DSR transfer curve) or $R_1 \approx 0.4\text{s}^{-1}$ (fold in the UNI transfer curve) in LessBias), indicating $R_1$-nulling.

### 4 The impact of curvature correction on $R_1$-maps

As the myelination of the cortex is influenced trivially by the local curvature, it makes sense to perform a correction for the local curvature, when sampling $R_1$-maps on the cortical surface. To illustrate that the effects of our $B_1^+$-manipulations are not affected by this, we show in Figure S3 both the curvature-corrected and uncorrected $R_1$-surfaces together with the estimated curvature. From the standard deviation surfaces it is seen that the variation in the estimated curvature across $B_1^+$-manipulation does not affect the $R_1$-maps.

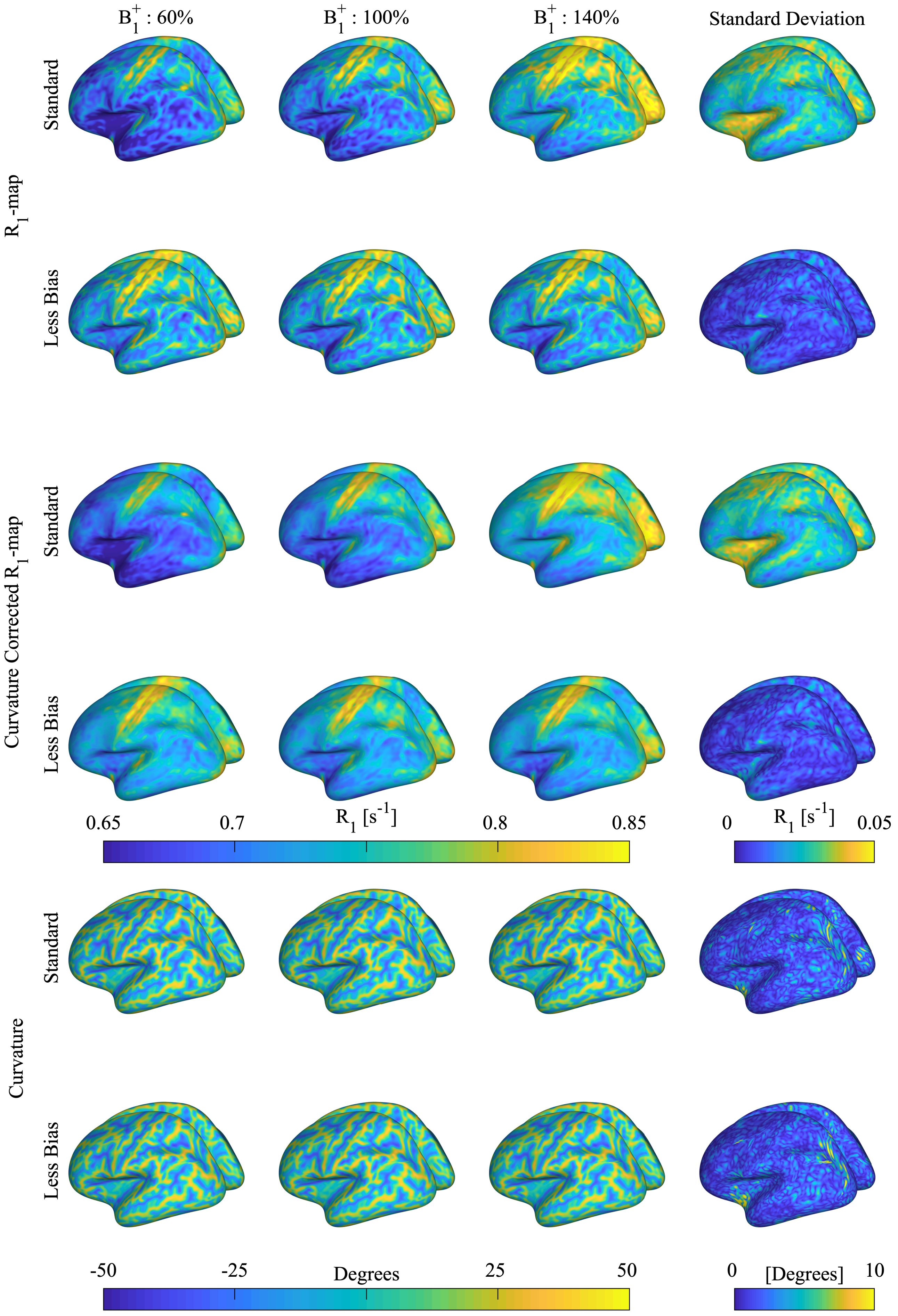


**Figure S3:** The effect of curvature correction on the estimated $R_1$-surfaces and the standard deviation across $B_1^+$ manipulations. By projecting out the local curvature of the $R_1$-values it is possible to remove curvature-induced variation in the $R_1$-mapping. As seen from the two bottom rows, the estimated curvatures are almost identical across protocols and $B_1^+$-manipulations. From the standard deviation of the curvature-uncorrected $R_1$-maps (two top rows) and curvature-corrected $R_1$-maps (two central rows), it is seen that these are entirely driven by the $B_1^+$-manipulations.

### 5 Robustness of CAT12 segmentation outputs

The volumes and thickness estimates outputted from the CAT12 segmentation was assessed towards robustness within and between sessions (Table S2), between subjects and $B_1^+$-manipulations (Table S3); and across sequence parameters )Table S4). From these tables it is clear that LessBias offers a substantial improvement in the consistency of the outputted tissue volumes, when $B_1^+$or sequence parameters are altered.

**Table S2:** Segmentation robustness within and across sessions. Top (Subject 1 Session 3) shows the variation of estimated volumes across repeated acquisitions of the two protocols. It is seen that LessBias shows a 2-3 fold larger standard deviation in estimated volumes, when compared to Standard, but in both cases the standard deviation across repetitions are for GM approximately 0.25 % of the corresponding volumes. Center (Subject 1) and bottom (Subject 2) show that Standard yields a 2.68 and 1.7 times higher standard deviation in the GM volume estimation than LessBias, respectively, when compared different sessions of the same subject.

| **Subject 1 Session 3** | Rep. | CSF [mL] | GM [mL] | WM [mL] | TIV [mL] | GM-thickness [mm] |
|---|---|---|---|---|---|---|
| Standard | 1 | 310.5 | 816.9 | 524.9 | 1652.3 | 2.34 |
| | 2 | 311.2 | 816.6 | 524.4 | 1652.2 | 2.31 |
| | 3 | 311.9 | 818.0 | 524.8 | 1654.7 | 2.31 |
| LessBias | 1 | 259.1 | 852.7 | 523.4 | 1635.1 | 2.40 |
| | 2 | 255.1 | 855.7 | 524.3 | 1635.1 | 2.40 |
| | 3 | 258.5 | 851.2 | 523.9 | 1633.6 | 2.44 |
| **Std(Standard)** | - | **0.7** | **0.7** | **0.3** | **1.4** | **0.01** |
| **Std(LessBias)** | - | **2.1** | **2.3** | **0.5** | **0.9** | **0.03** |

| **Subject 1** | Session | CSF [mL] | GM [mL] | WM [mL] | TIV [mL] | GM-thickness [mm] |
|---|---|---|---|---|---|---|
| Standard | 1 | 302.7 | 828.4 | 524.8 | 1655.9 | 2.31 |
| | 2 | 304.4 | 823.2 | 525.6 | 1653.2 | 2.34 |
| | 3 | 311.2 | 816.6 | 524.4 | 1652.2 | 2.31 |
| LessBias | 1 | 251.3 | 859.5 | 523.0 | 1633.8 | 2.43 |
| | 2 | 251.1 | 859.4 | 524.8 | 1635.3 | 2.41 |
| | 3 | 255.1 | 855.7 | 524.3 | 1635.1 | 2.40 |
| **Std(Standard)** | - | **4.5** | **5.9** | **0.6** | **1.9** | **0.01** |
| **Std(LessBias)** | - | **2.3** | **2.2** | **0.9** | **0.8** | **0.02** |

| **Subject 2** | Session | CSF [mL] | GM [mL] | WM [mL] | TIV [mL] | GM-thickness [mm] |
|---|---|---|---|---|---|---|
| Standard | 1 | 283.0 | 828.9 | 644.1 | 1756.1 | 2.41 |
| | 2 | 261.9 | 848.9 | 647.4 | 1758.3 | 2.38 |
| | 3 | 259.6 | 852.5 | 649.1 | 1761.1 | 2.46 |
| LessBias | 1 | 212.3 | 880.9 | 645.3 | 1738.5 | 2.36 |
| | 2 | 205.1 | 892.7 | 646.7 | 1744.5 | 2.35 |
| | 3 | 203.8 | 878.5 | 642.6 | 1724.9 | 2.58 |
| **Std(Standard)** | - | **12.9** | **12.7** | **2.5** | **2.5** | **0.04** |
| **Std(LessBias)** | - | **4.6** | **7.6** | **2.1** | **10.1** | **0.13** |

**Table S3:** Segmentation robustness across subjects. GM, WM, CSF and TIV volumes and GM average thickness for different protocols, subjects and $B_1^+$-manipulations from four sessions in three subjects. It is consistently seen that while Standard shows substantial variation in estimated volumes across $B_1^+$-modifications, LessBias is practically immune to the $B_1^+$-alterations. The standard deviation seen in the estimated volumes for LessBias is at the level of that seen in the top of Table S4 where each protocol was repeated three times without $B_1^+$-modifications. For Standard on the other hand the standard deviation across $B_1^+$-alterations is up to 22 times larger than what is seen for LessBias.

| **Subject 1 Session 1** | $B_1^+$ [%] | CSF [mL] | GM [mL] | WM [mL] | TIV [mL] | GM-thickness [mm] |
|---|---|---|---|---|---|---|
| Standard | 60 | 342.8 | 778.0 | 521.1 | 1642.0 | 2.16 |
| | 100 | 302.7 | 828.4 | 524.8 | 1655.9 | 2.31 |
| | 140 | 289.8 | 847.2 | 524.0 | 1661.0 | 2.38 |
| LessBias | 60 | 233.5 | 868.1 | 521.9 | 1623.5 | 2.56 |
| | 100 | 251.3 | 859.5 | 523.0 | 1633.8 | 2.43 |
| | 140 | 257.6 | 857.7 | 522.7 | 1638.0 | 2.40 |
| **Std(Standard)** | - | **27.7** | **35.8** | **1.9** | **9.8** | **0.11** |
| **Std(LessBias)** | - | **12.5** | **5.5** | **0.6** | **7.4** | **0.08** |

| **Subject 1 Session 2** | $B_1^+$ [%] | CSF [mL] | GM [mL] | WM [mL] | TIV [mL] | GM-thickness [mm] |
|---|---|---|---|---|---|---|
| Standard | 60 | 342.2 | 779.9 | 522.9 | 1645.0 | 2.20 |
| | 100 | 304.4 | 823.2 | 525.6 | 1653.2 | 2.34 |
| | 140 | 292.8 | 843.2 | 525.5 | 1661.5 | 2.38 |
| LessBias | 60 | 245.3 | 861.5 | 524.1 | 1630.9 | 2.45 |
| | 100 | 251.1 | 859.4 | 524.8 | 1635.3 | 2.41 |
| | 140 | 260.5 | 858.6 | 522.2 | 1641.3 | 2.31 |
| **Std(Standard)** | - | **25.9** | **32.4** | **1.5** | **8.2** | **0.09** |
| **Std(LessBias)** | - | **7.7** | **1.5** | **1.3** | **5.2** | **0.07** |

| **Subject 2 Session 2** | $B_1^+$ [%] | CSF [mL] | GM [mL] | WM [mL] | TIV [mL] | GM-thickness [mm] |
|---|---|---|---|---|---|---|
| Standard | 60 | 304.0 | 796.2 | 646.8 | 1747.0 | 2.28 |
| | 100 | 261.9 | 848.9 | 647.4 | 1758.3 | 2.38 |
| | 140 | 245.0 | 866.5 | 648.5 | 1760.0 | 2.43 |
| LessBias | 60 | 192.7 | 894.0 | 648.0 | 1734.7 | 2.48 |
| | 100 | 205.1 | 892.7 | 646.7 | 1744.5 | 2.35 |
| | 140 | 204.5 | 895.8 | 643.1 | 1743.4 | 2.37 |
| **Std(Standard)** | - | **30.4** | **36.6** | **0.9** | **7.1** | **0.08** |
| **Std(LessBias)** | - | **7.0** | **1.6** | **2.5** | **5.4** | **0.07** |

| **Subject 3 Session 1** | $B_1^+$ [%] | CSF [mL] | GM [mL] | WM [mL] | TIV [mL] | GM-thickness [mm] |
|---|---|---|---|---|---|---|
| Standard | 60 | 287.5 | 803.7 | 563.6 | 1654.8 | 2.32 |
| | 100 | 267.6 | 835.6 | 561.6 | 1664.7 | 2.44 |
| | 140 | 253.9 | 857.3 | 560.5 | 1671.7 | 2.62 |
| LessBias | 60 | 192.2 | 886.4 | 565.3 | 1643.9 | 2.59 |
| | 100 | 207.3 | 885.7 | 558.8 | 1651.8 | 2.56 |
| | 140 | 208.3 | 887.8 | 558.0 | 1654.1 | 2.52 |
| **Std(Standard)** | - | **16.9** | **27.0** | **1.6** | **8.5** | **0.15** |
| **Std(LessBias)** | - | **9.0** | **1.1** | **4.0** | **5.3** | **0.03** |

**Table S4:** Segmentation robustness across sequence parameters. Top (Subject 2 Session 1), it is seen that the estimated volumes for the two protocols change a lot when Standard is acquired with various number of slices, while this is not the case for LessBias. The standard deviation across slice modulations for Standard is 2-16 times greater than for LessBias, where the standard deviation is comparable to when no parameters are modified. In GM, the standard deviation of estimated GM volumes for Standard is almost 2% of the estimated volume. At the bottom (Subject 2 Session 3), the effect of $B_1^+$correction and presence of fat navigators (FatNavs) are shown. A • in the FatNavs column indicates that the acquisition was performed using the FatNavs-MP2RAGE sequence (Gallichan et al. MRM 75 (2016) 1030-1039), whereas a ◦ indicates that the product version without fat navigators was used. Both protocols were reconstructed on the scanner i.e. without performing the motion correction. For LessBias, the estimated volumes are practically unaffected by the fat navigators, while the standard deviation for Standard is 8 times larger than for LessBias. In GM, the standard deviation of estimated GM volumes for Standard is around 1% of the estimated volume. $B_1^+$-correction of Standard reduce the standard deviation between the protocols by a factor of 2.

| **Subject 2 Session 1** | #Slices | CSF [mL] | GM [mL] | WM [mL] | TIV [mL] | GM-thickness [mm] |
|---|---|---|---|---|---|---|
| Standard | 160 | 291.6 | 814.1 | 642.1 | 1747.9 | 2.40 |
| | 192 | 283.0 | 828.9 | 644.1 | 1756.1 | 2.41 |
| | 240 | 269.0 | 844.6 | 645.9 | 1759.5 | 2.45 |
| LessBias | 160 | 213.9 | 881.2 | 643.6 | 1738.7 | 2.39 |
| | 192 | 212.3 | 880.9 | 645.3 | 1738.5 | 2.36 |
| | 240 | 215.9 | 882.4 | 646.1 | 1744.4 | 2.37 |
| **Std(Standard)** | - | **11.4** | **15.2** | **1.9** | **5.9** | **0.03** |
| **Std(LessBias)** | - | **1.8** | **0.8** | **1.3** | **3.3** | **0.01** |

| **Subject 2 Session 3** | FatNavs | CSF [mL] | GM [mL] | WM [mL] | TIV [mL] | GM-Thickness [mm] |
|---|---|---|---|---|---|---|
| Standard | • | 259.6 | 852.5 | 649.1 | 1761.1 | 2.5 |
| | ◦ | 265.7 | 840.8 | 648.4 | 1754.8 | 2.5 |
| LessBias | • | 203.8 | 878.5 | 642.6 | 1724.9 | 2.6 |
| | ◦ | 203.0 | 877.0 | 642.6 | 1722.7 | 2.6 |
| **Std(Standard)** | - | **4.3** | **8.3** | **0.5** | **4.4** | **0.03** |
| **Std(LessBias)** | - | **0.6** | **1.0** | **0.0** | **1.6** | **0.01** |

### 6 Robustness of $R_1$-values

The resulting $R_1$-values (mean and standard deviation), from images segmented with CAT12, were assessed towards robustness within and across sessions (Table S5); across subjects and $B_1^+$-manipulations (Table S6); and sequence parameters, (Table S7). For each tissue type (GM, WM and CSF), a mask was created from the intersections between Standard and LessBias of the segmented volumes from the unmanipulated (B1+: 100%) Standard and LessBias images. In Figure S4, we show robustness of $R_1$-surfaces, togeher with 1D and 2D histograms for repeated acquisitions within the same session, and in Figure S5 and Figure S6 for repetition across multiple sessions. The surfaces in Figure S7 are the same as those found in Figure 4 of the main paper. From the 1D and 2D histograms, we can further appreciate the effects of the $B_1^+$-manipulations described in the paper. From the 1D histogram, it is confirmed that the LessBias protocol is indeed very robust to the $B_1^+$-manipulations, whereas the Standard protocol is very affected. In Figure S8, we confirm that the R1 estimates are robust to changes in the number of slices in the interval from 160 to 240. Finally in Figure S9, we confirm that our use of FatNavs are unlikely to have introduced any bias to our $R_1$-mapping.

**Table S5:** $R_1$-values (mean and standard deviation) within and across sessions.

| **Subject 1 Session 3** | Rep. | CSF [$s^{-1}$] | GM [$s^{-1}$] | WM [$s^{-1}$] |
|---|---|---|---|---|
| Standard | 1 | 0.401 ± 0.170 | 0.712 ± 0.079 | 1.186 ± 0.077 |
| | 2 | 0.400 ± 0.173 | 0.712 ± 0.077 | 1.188 ± 0.079 |
| | 3 | 0.402 ± 0.176 | 0.712 ± 0.079 | 1.184 ± 0.078 |
| LessBias | 1 | 0.328 ± 0.265 | 0.732 ± 0.084 | 1.228 ± 0.089 |
| | 2 | 0.333 ± 0.272 | 0.734 ± 0.088 | 1.226 ± 0.090 |
| | 3 | 0.335 ± 0.274 | 0.734 ± 0.087 | 1.225 ± 0.089 |
| **Standard: Mean ± Std** | - | **0.401 ± 0.001** | **0.712 ± 0.000** | **1.186 ± 0.002** |
| **LessBias: Mean ± Std** | - | **0.332 ± 0.004** | **0.733 ± 0.001** | **1.226 ± 0.002** |

| **Subject 1** | Session | CSF [$s^{-1}$] | GM [$s^{-1}$] | WM [$s^{-1}$] |
|---|---|---|---|---|
| Standard | 1 | 0.398 ± 0.170 | 0.710 ± 0.077 | 1.186 ± 0.078 |
| | 2 | 0.400 ± 0.180 | 0.710 ± 0.077 | 1.188 ± 0.078 |
| | 3 | 0.401 ± 0.178 | 0.711 ± 0.077 | 1.188 ± 0.079 |
| LessBias | 1 | 0.324 ± 0.263 | 0.731 ± 0.084 | 1.228 ± 0.087 |
| | 2 | 0.329 ± 0.278 | 0.729 ± 0.084 | 1.225 ± 0.086 |
| | 3 | 0.328 ± 0.273 | 0.732 ± 0.083 | 1.228 ± 0.088 |
| **Standard: Mean ± Std** | - | **0.400 ± 0.002** | **0.710 ± 0.001** | **1.187 ± 0.001** |
| **LessBias: Mean ± Std** | - | **0.327 ± 0.003** | **0.731 ± 0.002** | **1.227 ± 0.002** |

| **Subject 2** | Session | CSF [$s^{-1}$] | GM [$s^{-1}$] | WM [$s^{-1}$] |
|---|---|---|---|---|
| Standard | 1 | 0.439 ± 0.209 | 0.713 ± 0.086 | 1.219 ± 0.081 |
| | 2 | 0.453 ± 0.239 | 0.705 ± 0.084 | 1.207 ± 0.077 |
| | 3 | 0.423 ± 0.179 | 0.711 ± 0.080 | 1.208 ± 0.072 |
| LessBias | 1 | 0.380 ± 0.327 | 0.733 ± 0.089 | 1.262 ± 0.093 |
| | 2 | 0.399 ± 0.347 | 0.725 ± 0.090 | 1.251 ± 0.087 |
| | 3 | 0.353 ± 0.253 | 0.727 ± 0.082 | 1.250 ± 0.080 |
| **Standard: Mean ± Std** | - | **0.438 ± 0.015** | **0.710 ± 0.004** | **1.211 ± 0.006** |
| **LessBias: Mean ± Std** | - | **0.378 ± 0.023** | **0.728 ± 0.004** | **1.254 ± 0.007** |

**Table S6:** $R_1$-values (mean and standard deviation) across subjects. In all four sessions Standard and LessBias were subject to $B_1^+$manipulations through changes of the FA (60, 100, and 140 %), see Sec. 2. We see a substantially smaller $R_1$standard deviation for LessBias in GM and WM than for Standard. Improvements range from a factor of 3.3 to 21.5. For CSF, LessBias ranges from 2 to 1.04 times higher than that of Standard.

| **Subject 1 Session 1** | $B_1^+$ [%] | CSF [s$^{-1}$] | GM [s$^{-1}$] | WM [s$^{-1}$] |
|---|---|---|---|---|
| Standard | 60 | 0.432 ± 0.145 | 0.681 ± 0.078 | 1.135 ± 0.085 |
| | 100 | 0.391 ± 0.164 | 0.709 ± 0.077 | 1.186 ± 0.078 |
| | 140 | 0.404 ± 0.179 | 0.750 ± 0.084 | 1.254 ± 0.079 |
| LessBias | 60 | 0.313 ± 0.328 | 0.742 ± 0.088 | 1.219 ± 0.098 |
| | 100 | 0.318 ± 0.256 | 0.731 ± 0.087 | 1.227 ± 0.090 |
| | 140 | 0.353 ± 0.250 | 0.732 ± 0.086 | 1.251 ± 0.090 |
| **Standard: Mean ± Std** | - | **0.409 ± 0.021** | **0.713 ± 0.035** | **1.192 ± 0.060** |
| **LessBias: Mean ± Std** | - | **0.328 ± 0.022** | **0.735 ± 0.006** | **1.232 ± 0.017** |

| **Subject 1 Session 2** | $B_1^+$ [%] | CSF [s$^{-1}$] | GM [s$^{-1}$] | WM [s$^{-1}$] |
|---|---|---|---|---|
| Standard | 60 | 0.427 ± 0.157 | 0.686 ± 0.078 | 1.136 ± 0.082 |
| | 100 | 0.397 ± 0.176 | 0.710 ± 0.077 | 1.187 ± 0.079 |
| | 140 | 0.420 ± 0.191 | 0.768 ± 0.087 | 1.285 ± 0.084 |
| LessBias | 60 | 0.311 ± 0.314 | 0.737 ± 0.088 | 1.216 ± 0.095 |
| | 100 | 0.327 ± 0.272 | 0.730 ± 0.088 | 1.223 ± 0.088 |
| | 140 | 0.372 ± 0.265 | 0.736 ± 0.088 | 1.259 ± 0.092 |
| **Standard: Mean ± Std** | - | **0.415 ± 0.016** | **0.721 ± 0.042** | **1.203 ± 0.076** |
| **LessBias: Mean ± Std** | - | **0.337 ± 0.032** | **0.734 ± 0.004** | **1.233 ± 0.023** |

| **Subject 2 Session 2** | $B_1^+$ [%] | CSF [s$^{-1}$] | GM [s$^{-1}$] | WM [s$^{-1}$] |
|---|---|---|---|---|
| Standard | 60 | 0.468 ± 0.192 | 0.678 ± 0.088 | 1.150 ± 0.087 |
| | 100 | 0.435 ± 0.223 | 0.706 ± 0.085 | 1.207 ± 0.077 |
| | 140 | 0.453 ± 0.228 | 0.745 ± 0.093 | 1.269 ± 0.083 |
| LessBias | 60 | 0.376 ± 0.387 | 0.734 ± 0.096 | 1.238 ± 0.101 |
| | 100 | 0.380 ± 0.328 | 0.727 ± 0.096 | 1.249 ± 0.090 |
| | 140 | 0.415 ± 0.315 | 0.731 ± 0.094 | 1.272 ± 0.095 |
| **Standard: Mean ± Std** | - | **0.452 ± 0.017** | **0.709 ± 0.034** | **1.209 ± 0.059** |
| **LessBias: Mean ± Std** | - | **0.390 ± 0.021** | **0.731 ± 0.004** | **1.253 ± 0.017** |

| **Subject 3 Session 1** | $B_1^+$ [%] | CSF [s$^{-1}$] | GM [s$^{-1}$] | WM [s$^{-1}$] |
|---|---|---|---|---|
| Standard | 60 | 0.446 ± 0.162 | 0.669 ± 0.075 | 1.100 ± 0.076 |
| | 100 | 0.416 ± 0.181 | 0.695 ± 0.075 | 1.147 ± 0.073 |
| | 140 | 0.440 ± 0.194 | 0.753 ± 0.085 | 1.243 ± 0.081 |
| LessBias | 60 | 0.341 ± 0.344 | 0.719 ± 0.080 | 1.173 ± 0.087 |
| | 100 | 0.344 ± 0.290 | 0.715 ± 0.079 | 1.182 ± 0.082 |
| | 140 | 0.384 ± 0.272 | 0.715 ± 0.077 | 1.209 ± 0.084 |
| **Standard: Mean ± Std** | - | **0.434 ± 0.016** | **0.706 ± 0.043** | **1.163 ± 0.073** |
| **LessBias: Mean ± Std** | - | **0.356 ± 0.024** | **0.716 ± 0.002** | **1.188 ± 0.019** |

**Table S7:** $R_1$-values (mean and standard deviation) across sequence parameters. In the top (Subject 2 Session 1), the number of slices were changed. In the bottom (Subject 2 Session 3) a • in the FatNavs column indicates that the acquisition was performed using the FatNavs-MP2RAGE sequence (Gallichan et al. MRM 75 (2016) 1030-1039), whereas a ◦ indicates that the product version without fat navigators was used. Both sequences were reconstructed on the scanner i.e. without performing the motion correction. All scans were obtained with the same 100 % FA setting, and we see that the standard deviations for all tissue types are below 0.007 $s^{-1}$ for both Standard and LessBias.

| **Subject 2 Session 1** | #Slices | CSF [$s^{-1}$] | GM [$s^{-1}$] | WM [$s^{-1}$] |
|---|---|---|---|---|
| Standard | 160 | 0.457 ± 0.206 | 0.714 ± 0.084 | 1.216 ± 0.082 |
| | 192 | 0.439 ± 0.209 | 0.713 ± 0.086 | 1.219 ± 0.081 |
| | 240 | 0.429 ± 0.232 | 0.712 ± 0.088 | 1.219 ± 0.081 |
| LessBias | 160 | 0.381 ± 0.317 | 0.738 ± 0.088 | 1.267 ± 0.092 |
| | 192 | 0.380 ± 0.327 | 0.733 ± 0.089 | 1.262 ± 0.093 |
| | 240 | 0.394 ± 0.351 | 0.730 ± 0.095 | 1.263 ± 0.092 |
| **Standard: Mean ± Std** | - | **0.442 ± 0.014** | **0.713 ± 0.001** | **1.218 ± 0.002** |
| **LessBias: Mean ± Std** | - | **0.385 ± 0.008** | **0.734 ± 0.004** | **1.264 ± 0.003** |

| **Subject 2 Session 3** | FatNavs | CSF [$s^{-1}$] | GM [$s^{-1}$] | WM [$s^{-1}$] |
|---|---|---|---|---|
| Standard | • | 0.423 ± 0.179 | 0.711 ± 0.080 | 1.208 ± 0.072 |
| | ◦ | 0.425 ± 0.178 | 0.703 ± 0.082 | 1.199 ± 0.074 |
| LessBias | • | 0.353 ± 0.253 | 0.727 ± 0.082 | 1.250 ± 0.080 |
| | ◦ | 0.359 ± 0.251 | 0.725 ± 0.085 | 1.243 ± 0.083 |
| **Standard: Mean ± Std** | - | **0.424 ± 0.001** | **0.707 ± 0.006** | **1.204 ± 0.007** |
| **LessBias: Mean ± Std** | - | **0.356 ± 0.004** | **0.726 ± 0.002** | **1.246 ± 0.006** |

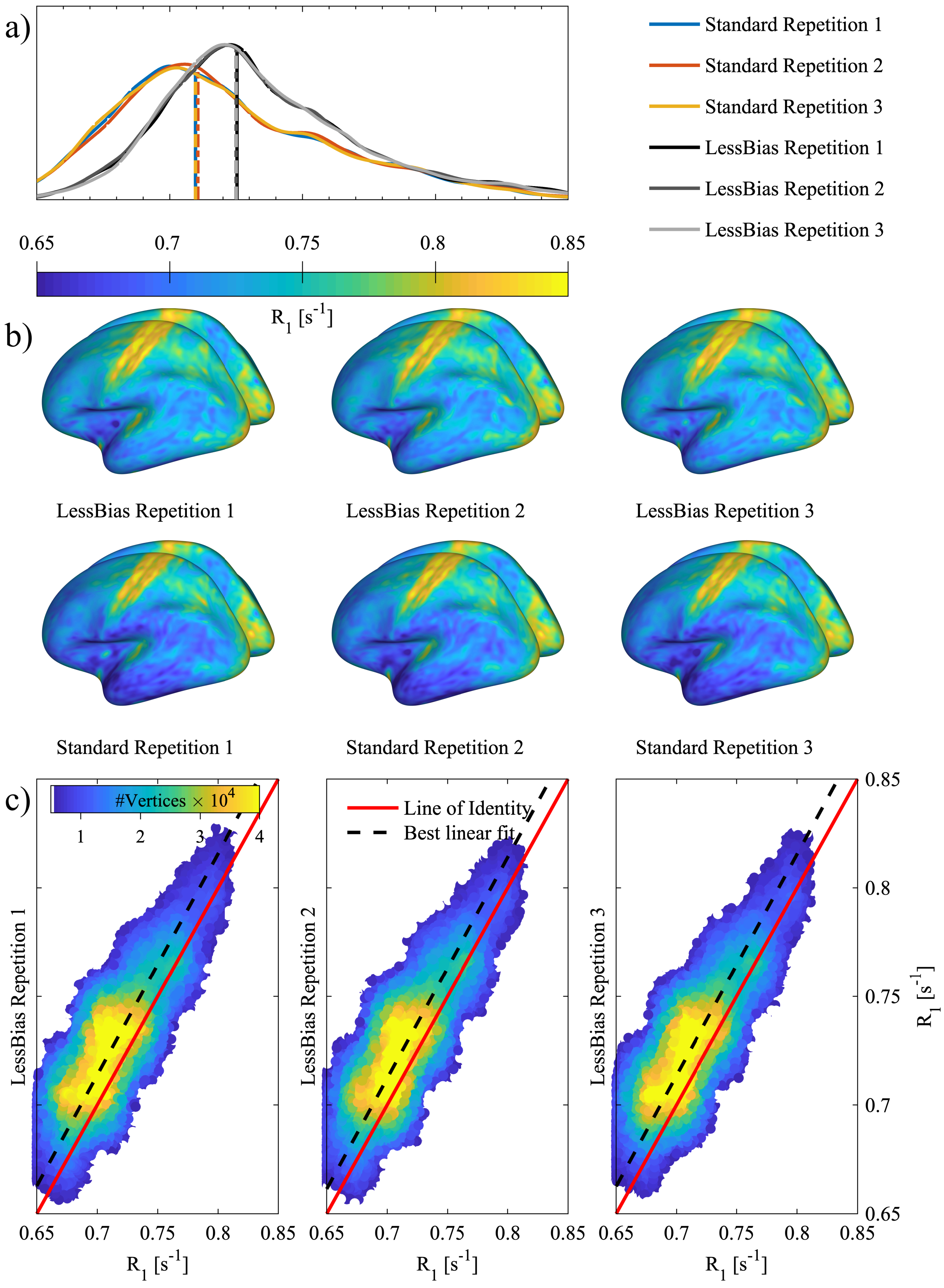


**Figure S4:** GM $R_1$-assessment, Subject 1 Session 3, regarding multiple repetitions. Panel a) shows distributions of $R_1$-values on the central GM-surface for both Standard and LessBias, across three repetitions within the same scanning session. In panel b) the surfaces corresponding to the curves in panel a) are shown. In panel c) 2D histograms show the relationship between Standard and LessBias central, cortical GM $R_1$-values. The red lines indicate the perfect relationship between the data and the dashed, black lines indicate the best linear fit with an intercept of zero.

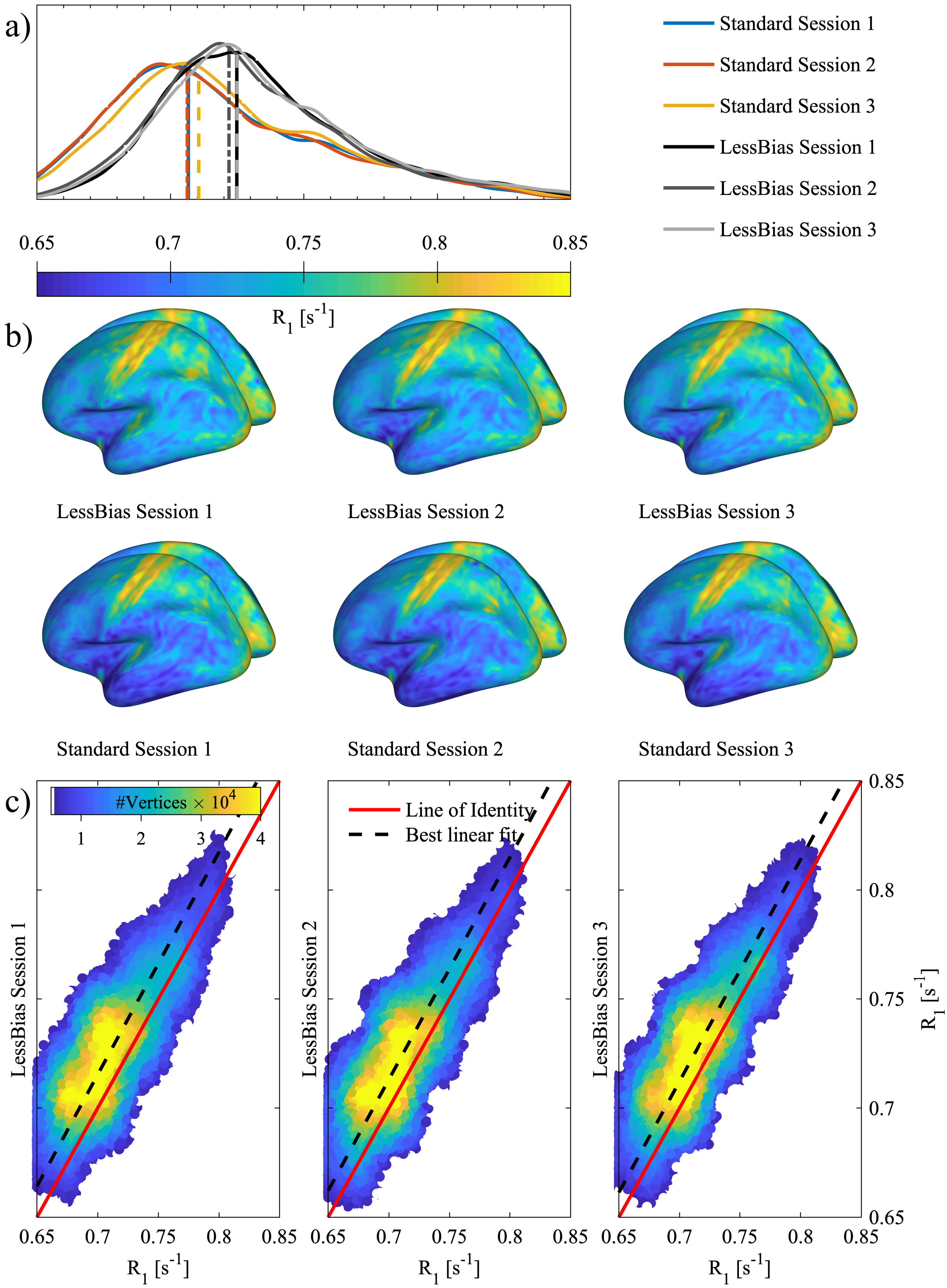


**Figure S5:** GM $R_1$-assessment, Subject 1 Session 1-3, regarding multiple sessions. Panel a) shows distributions of $R_1$-values on the central GM-surface for both Standard and LessBias, across three scanning sessions. In panel b) the surfaces corresponding to the curves in panel a) are shown. In panel c) 2D histograms show the relationship between the Standard and the LessBias central, cortical GM $R_1$-values. The red lines indicate the perfect relationship between the data and the dashed, black lines indicate the best linear fit with an intercept of zero.

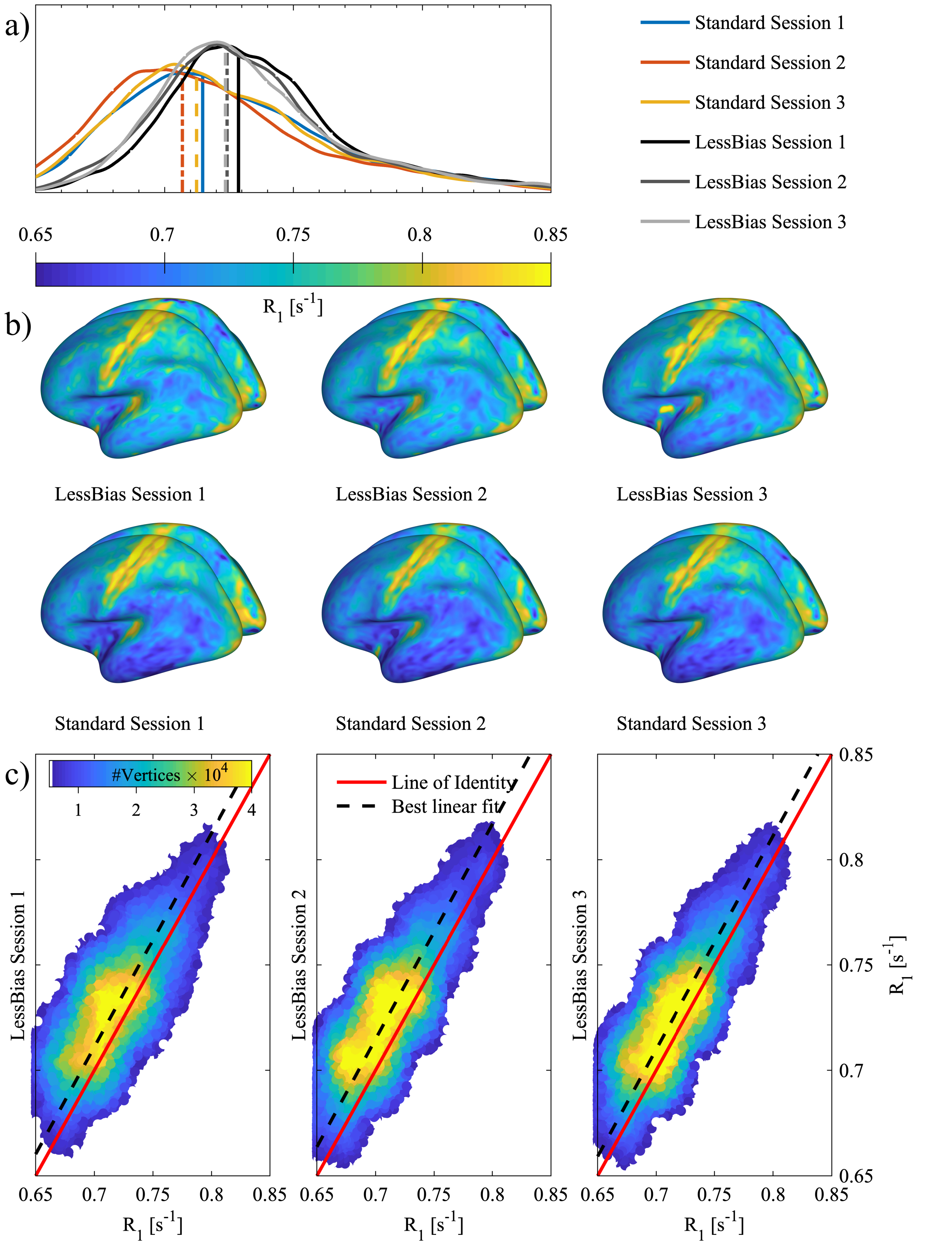


**Figure S6:** GM $R_1$-assessment, Subject 2 Session 1-3, regarding multiple sessions. Panel a) shows the distribution of $R_1$-values on the central GM-surface for both Standard and LessBias across three scanning sessions. Panel b) shows the surfaces corresponding to the curves in panel a). Panel c) shows 2D histograms of the relationship between Standard and LessBias central GM cortical $R_1$-values. It is seen that both Standard and LessBias are quite consistent across sessions, although not quite at the level seen in subject1 Figure S5 where a less motion was present. See Table S5 for a whole brain comparison.

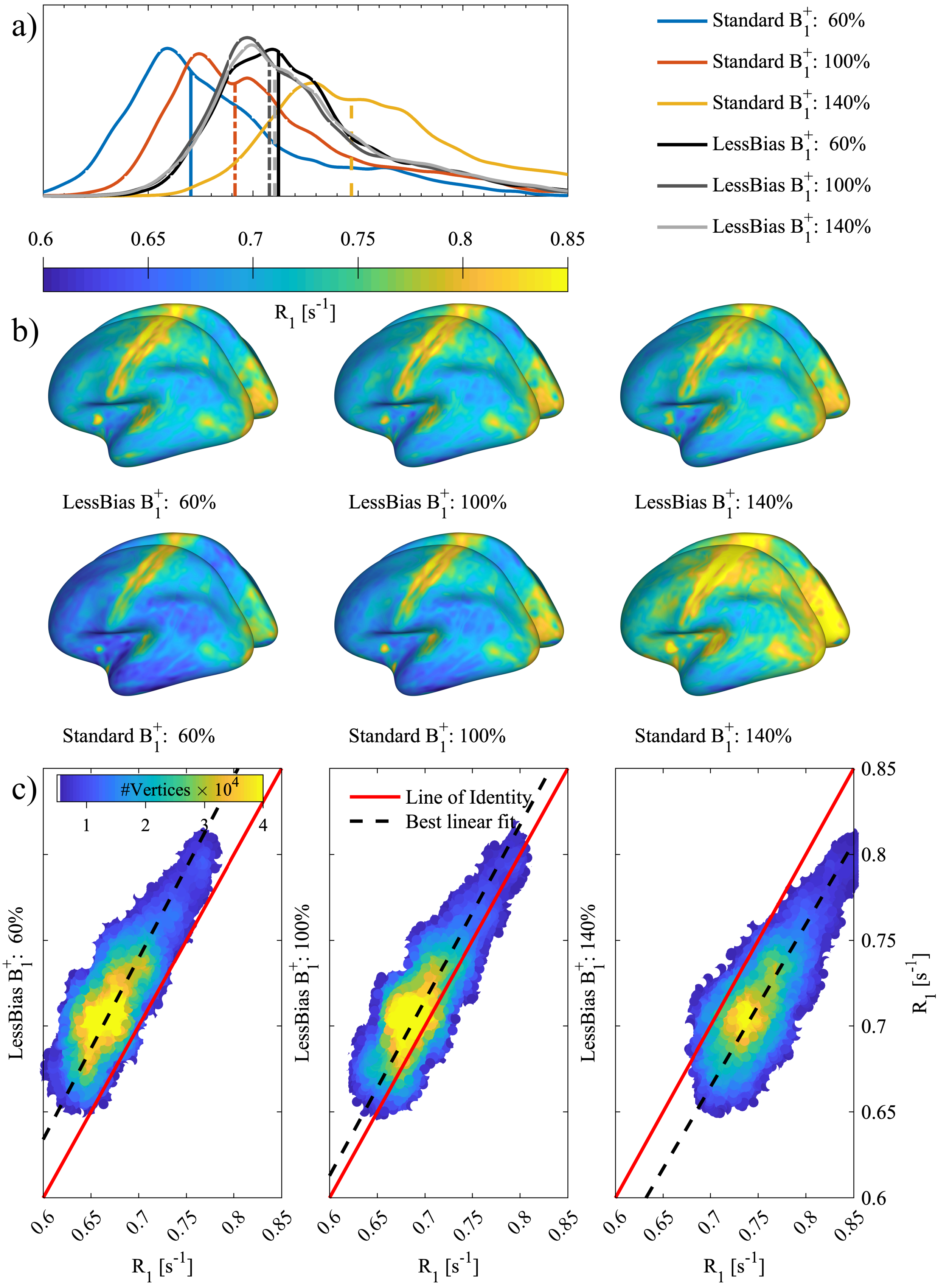


**Figure S7:** GM $R_1$-assessment, Subject 3 Session 1, regarding multiple $B_1^+$-values. Panel a) shows distributions of $R_1$-values on the central GM-surface for both Standard and LessBias, across three $B_1^+$-manipulations. From the distributions and their median values (indicated with vertical lines) it is clear that LessBias method is quite consistent across $B_1^+$-values, but Standard fluctuates rapidly. Panel b) shows the surfaces corresponding to the curves in panel a). The Standard also visually shows a substantial fluctuation in $R_1$-values, whereas LessBias shows much less variation. Panel c) shows 2D histograms of the relationship between Standard and LessBias central GM cortical $R_1$-values. It is overall seen that LessBias presents with more constant distributions across the $B_1^+$-manipulations than Standard, and that Standard agrees most with LessBias when $B_1^+$: 100 %.

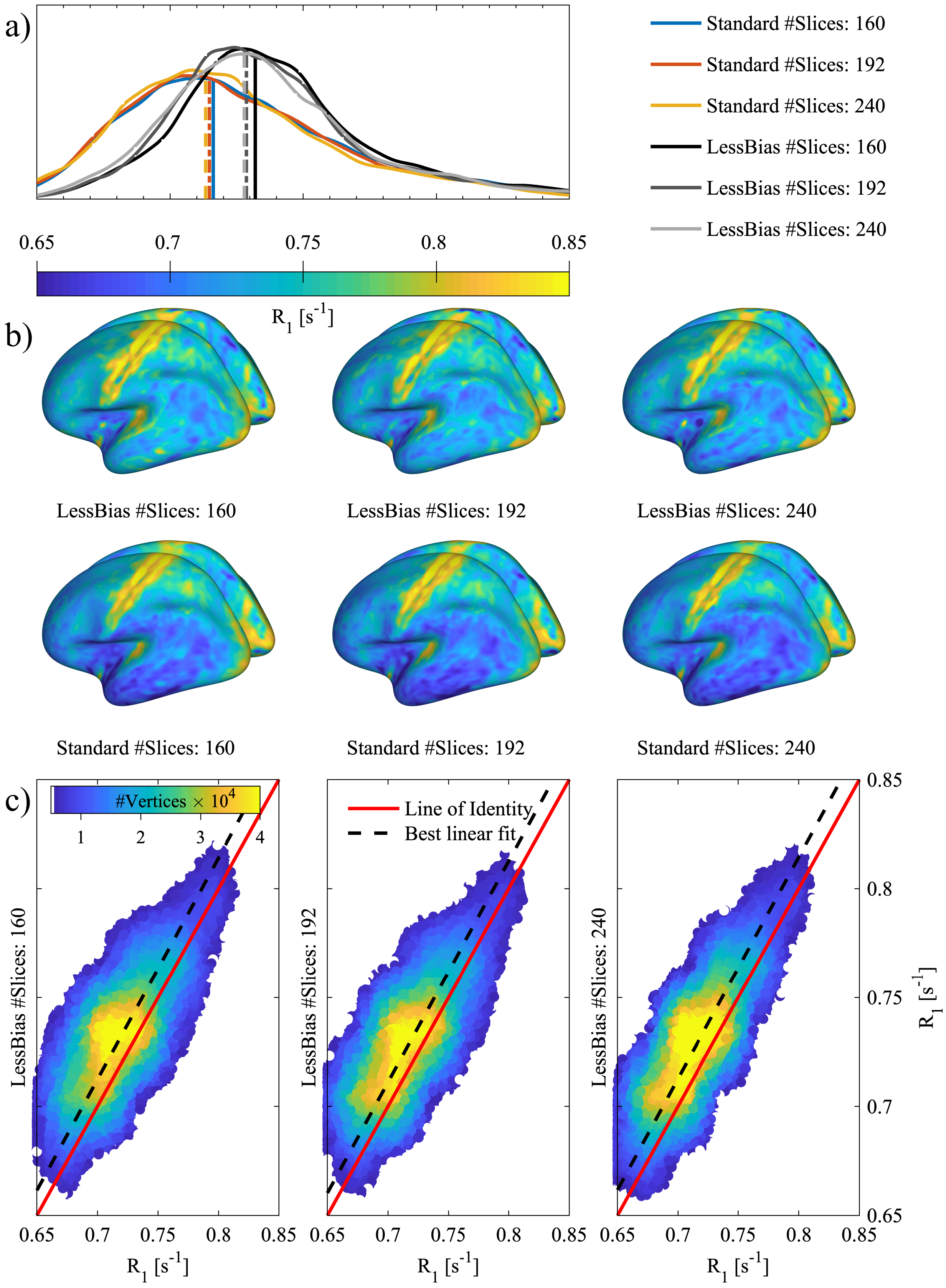


**Figure S8:** GM $R_1$-assessment, Subject 2 Session 1, regarding number of slices. Panel a) shows distributions of $R_1$-values on the central GM-surface for both Standard and LessBias for the different number of slices. Panel b) shows the surfaces corresponding to the curves in panel a). Panel c) shows 2D histograms of the relationship between Standard and LessBias central GM cortical $R_1$-values. It is seen that in the examined interval, both protocols seem quite robust to changes in the number of slices. See Table S7 for a whole brain comparison.

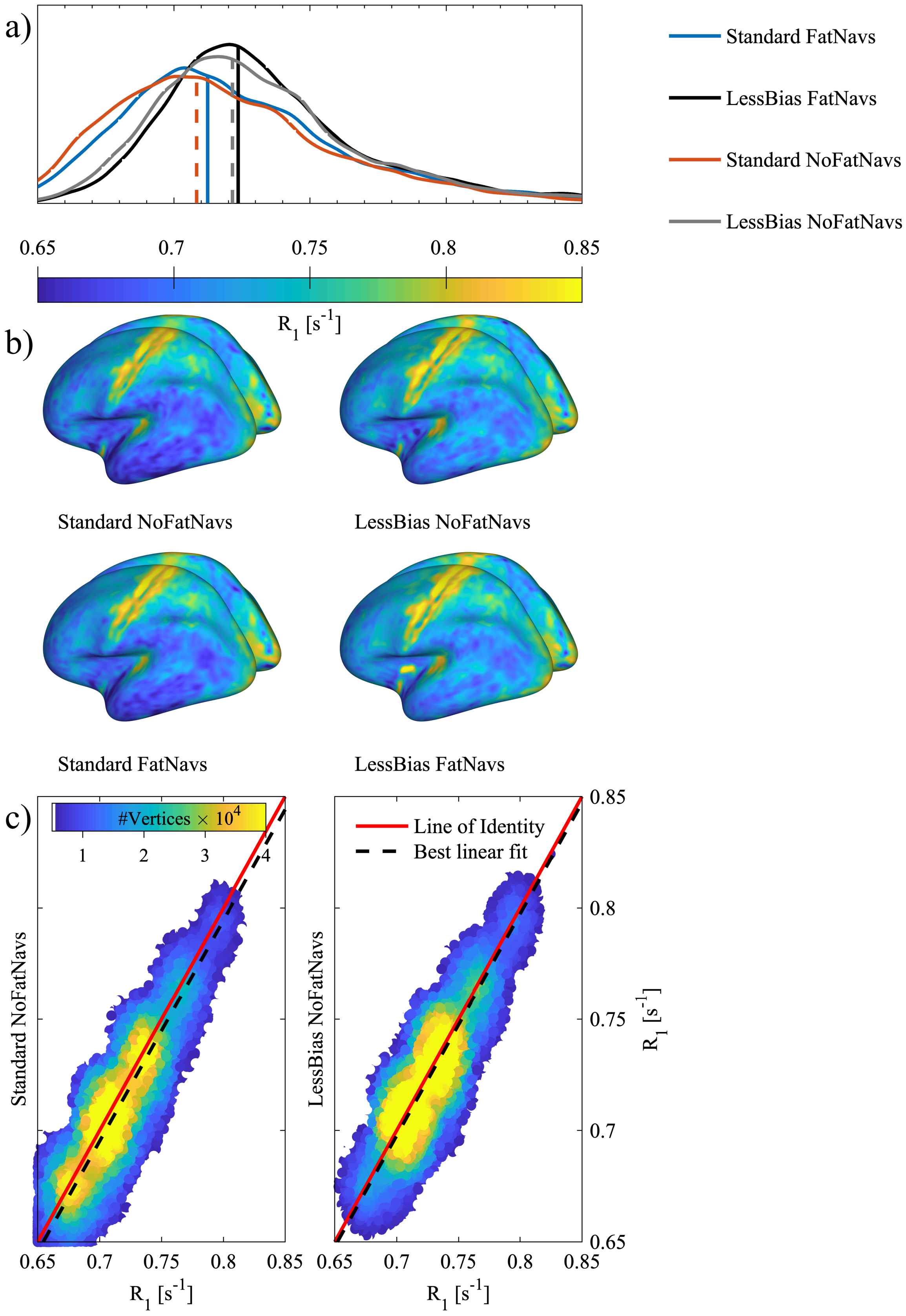


**Figure S9:** GM $R_1$-assessment, Subject 2 Session 3, regarding the fat navigator module. Panel a) shows distributions of $R_1$-values on the central GM-surface for both Standard and LessBias, with and without a fat navigator module present in the sequence. To be clear, "FatNavs" indicates that the acquisition was performed using the FatNavs-MP2RAGE sequence (Gallichan et al. MRM 75 (2016) 1030-1039), whereas "NoFatNavs" indicates that the MP2RAGE product version without fat navigators was used. The data in this analysis from the "FatNavs" scans, however, was the non-motion corrected data set. From the distributions (median values indicated with vertical lines) it is clear that the fat navigator pulse have no effect by itself. Panel b) shows the surfaces corresponding to the curves in panel a). In panel c), 2D histograms also show the lack of effect from the fat navigator module. The red lines indicate the perfect relationship between the data and the dashed, black lines indicate the best linear fit with an intercept of zero. An almost perfect correlation between the two MP2RAGE protocols is seen for both LessBias and Standard, indicating that the FatNavs pulse did not, by it self, influence the estimated $R_1$-values.

### 7 Effect of $B_1^+$-correction.

While the un-manipulated $R_1$-values from Standard and LessBias are fairly similar, there are reasons to believe that the un-manipulated Standard $R_1$-map is already affected by $B_1^+$-inhomogeneity. To compare our proposed method to a $B_1^+$-map-based correction method, we followed the SA2RAGE $B_1^+$-correction method (Eggenschwiler et al. MRM 67 (2012) 1609-1619) and acquired such scans accordingly. However, as the genuine SA2RAGE sequence was not available to us at time for this study, we used our MP2RAGE product sequence and switched the MP pulse to saturation mode instead of inversion. As a consequence hereof we do not directly get the real-value-ratio image, which was used for the $B_1^+$-mapping by Eggenschwiler et al. To accommodate this challenge we reconstructed the raw data to both produce an image containing the ratio between the real values of the two saturation images, but also a unified image, as outputted by the modified MP2RAGE sequence on the scanner. By modifying the equation behind the SA2RAGE LUT, we could convert both unified RAGE images and real-value-ratio images into $B_1^+$-maps. As seen in see Figure S10, we found the modified approach substantially more stable, and hence we used unified RAGE images for our in-vivo $B_1^+$-corrections.

In order to be able to perform a $B_1^+$-correction of protocols with a non-bijective transfer curve, we developed a 3D-LUT $B_1^+$-correction, which can work on $I_{\text{UNI}}$, $I_{\text{DSR}}$ and $B_1^+$-images simultaneously. In Figure S11 to Figure S15, we show the impact of $B_1^+$-correction of the images from the Standard protocol. For these corrections we used a 3D-LUT weighting of 100:0:100 thereby ignoring the information in $I_{\text{DSR}}$ for the Standard protocol. As an example of a true 3D-LUT $B_1^+$-correction, we show in Figure S16 the result of $B_1^+$-correction of a LessBias acquisition with 240 slices. In this case, we used a weighting of 100:1:100 to also incorporate the information found in $I_{\text{DSR}}$. Even with a protocol slightly more sensitive to $B_1^+$-inhomogenieties than the one acquired with 192 slices, we see almost no effect of the $B_1^+$-correction of the images from the LessBias acquisition.

Of note is that the same principles by which the 3D-LUT is performed, can trivially be extended to an ND-LUT application. Additional information, like $M_0$-maps, may be provided for further corrections, at the cost of computation time. We speculate that it may further be possible to filter out and estimate maps for one type of artefact outside of the provided information, by using the distance metric provided by the ND-LUT to determine the optimal solution across the estimated value, but this is subject of future investigation.

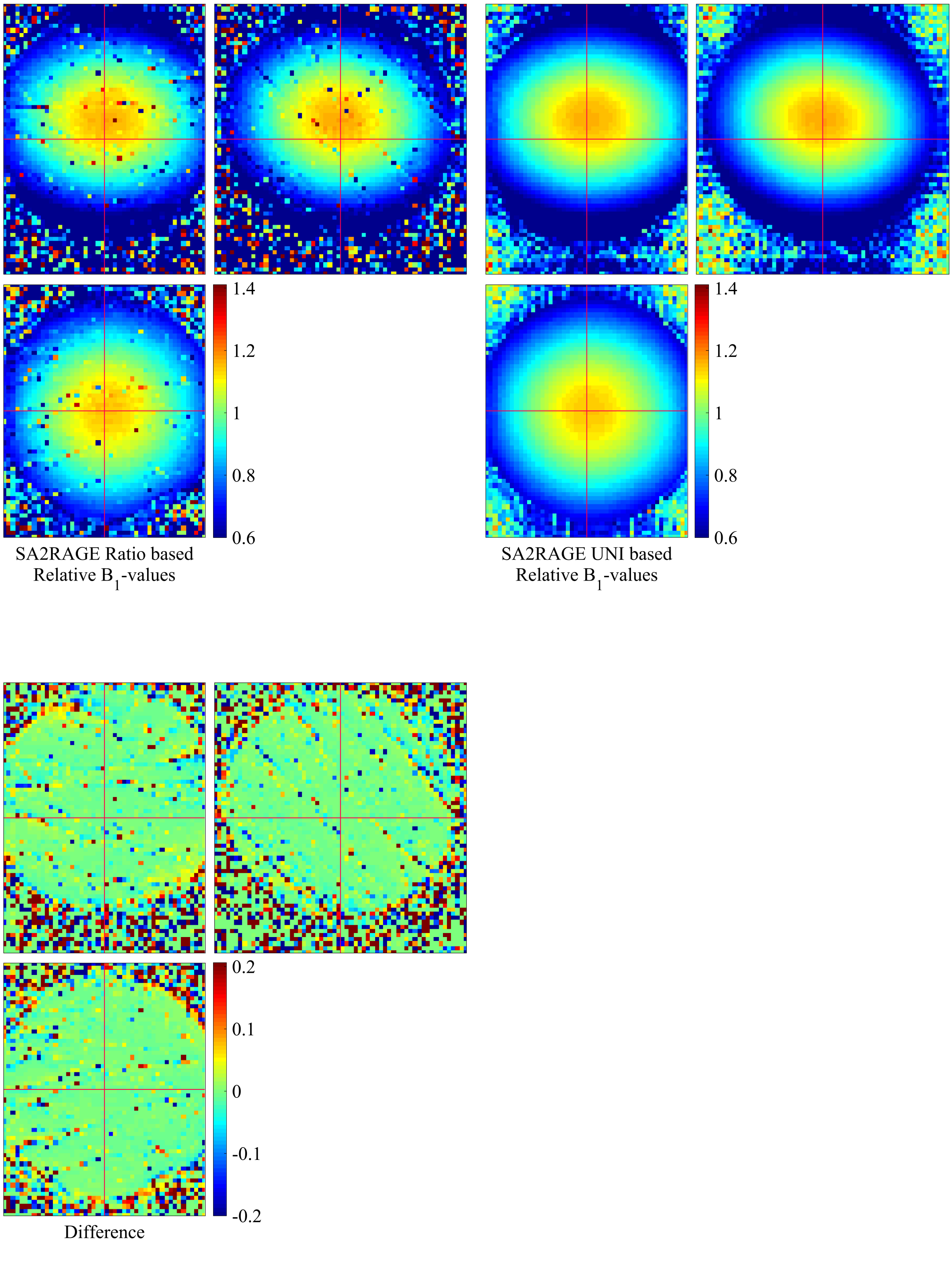


**Figure S10:** Phantom $B_1^+$-maps. Instead of the original, ratio-based SA2RAGE method (three, top left images), we estimated our $B_1^+$-maps (three, top right images) through the $I_{\text{UNI}}$-values and performed the correct via a 3D-LUT procedure. The essential difference between the two procedures is shown in the three bottom left images.

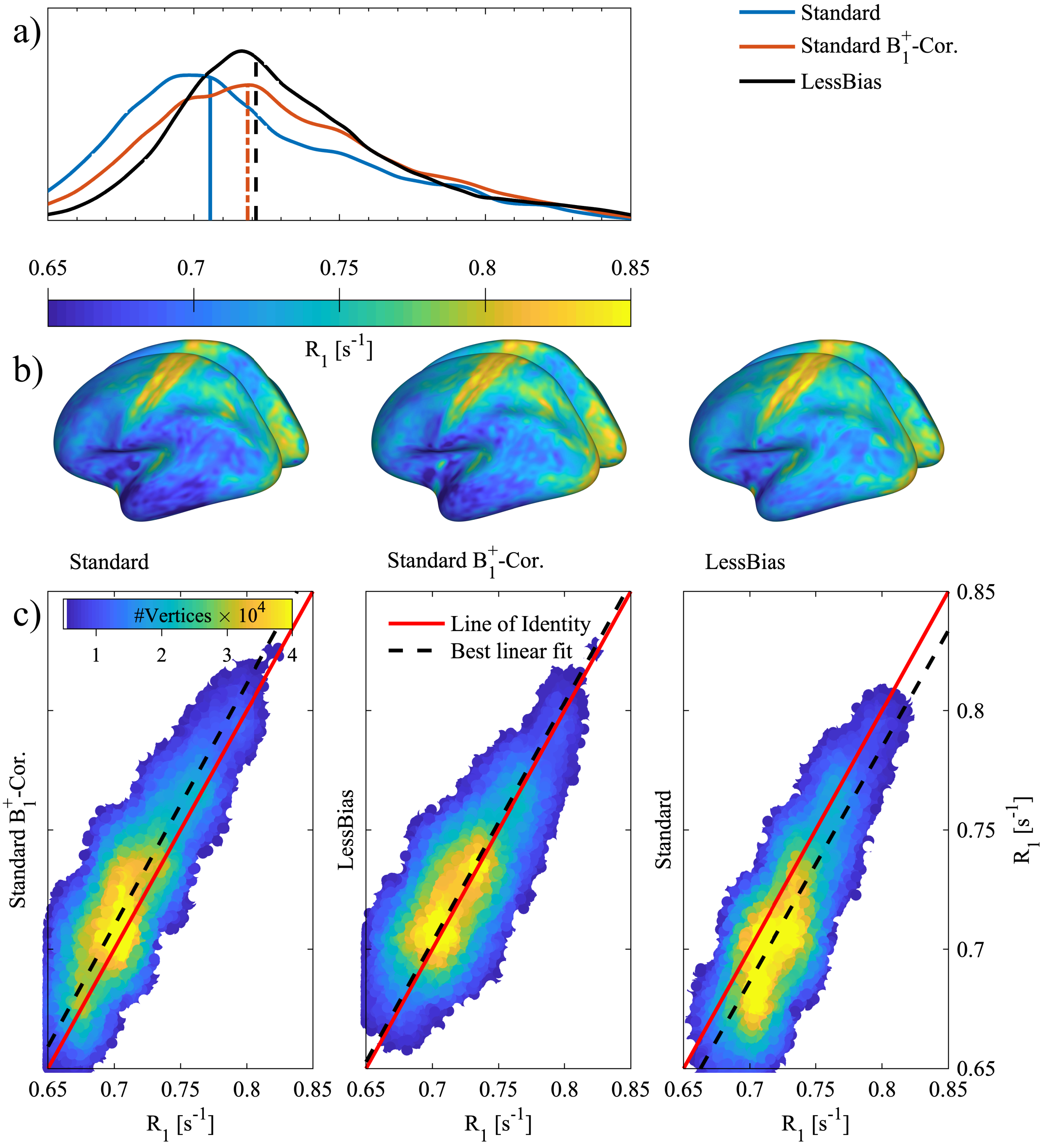


**Figure S11:** GM $R_1$-assessment, Subject 1 Session 2, regarding $B_1^+$-correction. Panel a) shows the distribution of $R_1$-values on the central GM-surface for both Standard and LessBias, As well as for a $B_1^+$-corrected version of Standard. From the distributions and their median values (indicated with vertical lines) it is clear that the $R_1$-values from Standard resemble those of LessBias, when the $R_1$-map from Standard is $B_1^+$-corrected. Panel b) shows the surfaces corresponding to the curves in panel a). A shift towards higher $R_1$-values at the cortex is seen, when the $R_1$-map from Standard is $B_1^+$-corrected. Panel c) shows 2D histograms for the three possible combinations of the maps in panel b). The red lines indicate the perfect relationship between the data and the dashed, black lines indicate the best linear fit with an intercept of zero. It is seen from the plots that while the maps from LessBias and Standard are fairly similar, the best correspondence is found between the $R_1$-maps from $B_1^+$-corrected Standard and LessBias.

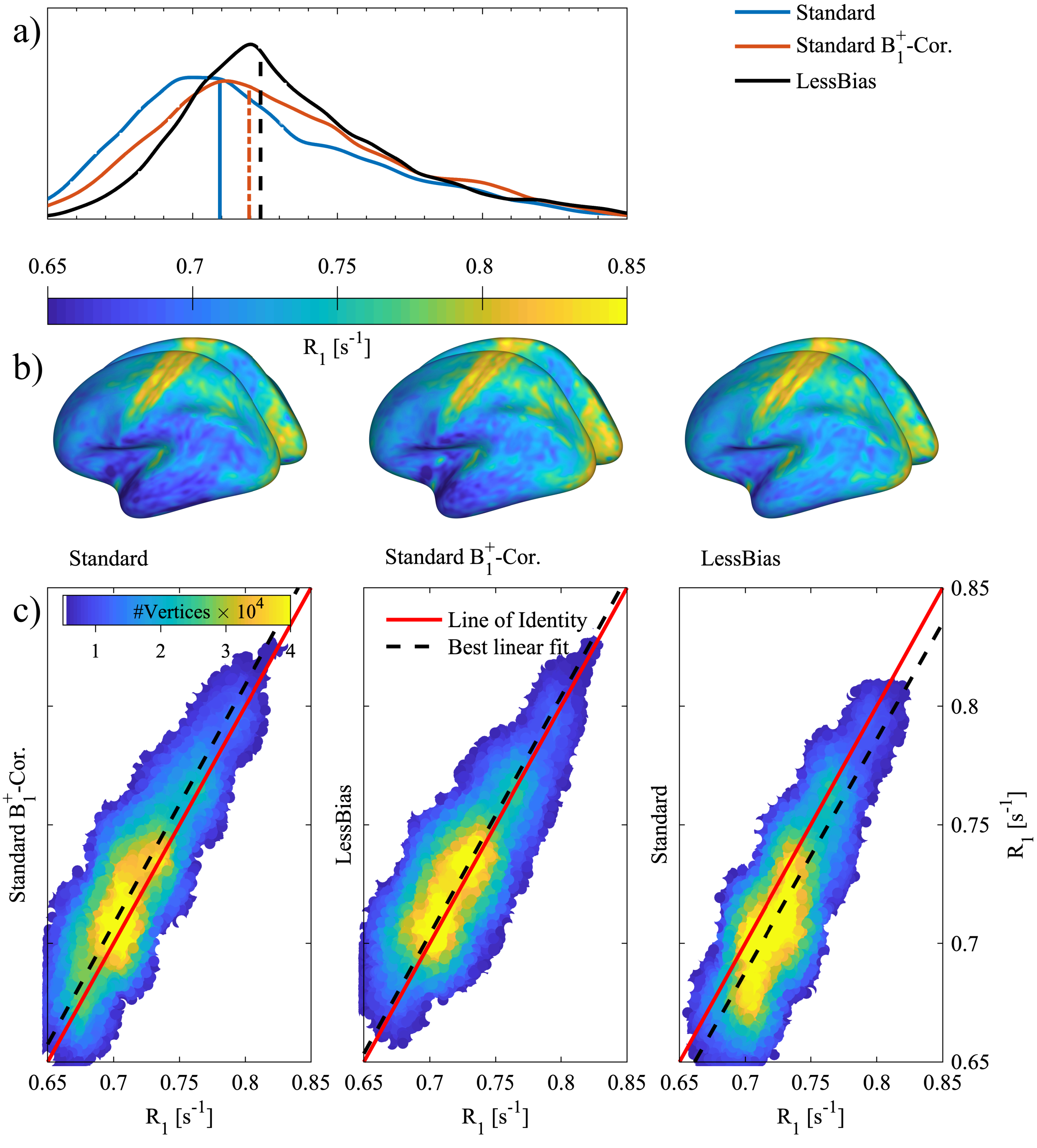


**Figure S12:** GM $R_1$-assessment, Subject 1 Session 3, regarding $B_1^+$-correction. Panel a) shows the distribution of $R_1$-values on the central GM-surface for both Standard and LessBias, As well as for a $B_1^+$-corrected version of Standard. From the distributions and their median values (indicated with vertical lines) it is clear that the $R_1$-values from Standard resemble those of LessBias, when the $R_1$-map from Standard is $B_1^+$-corrected. Panel b) shows the surfaces corresponding to the curves in panel a). A shift towards higher $R_1$-values at the cortex is seen, when the $R_1$-map from Standard is $B_1^+$-corrected. Panel c) shows 2D histograms for the three possible combinations of the maps in panel b). The red lines indicate the perfect relationship between the data and the dashed, black lines indicate the best linear fit with an intercept of zero. It is seen from the plots that while the maps from LessBias and Standard are fairly similar, the best correspondence is found between the $R_1$-maps from $B_1^+$-corrected Standard and LessBias.

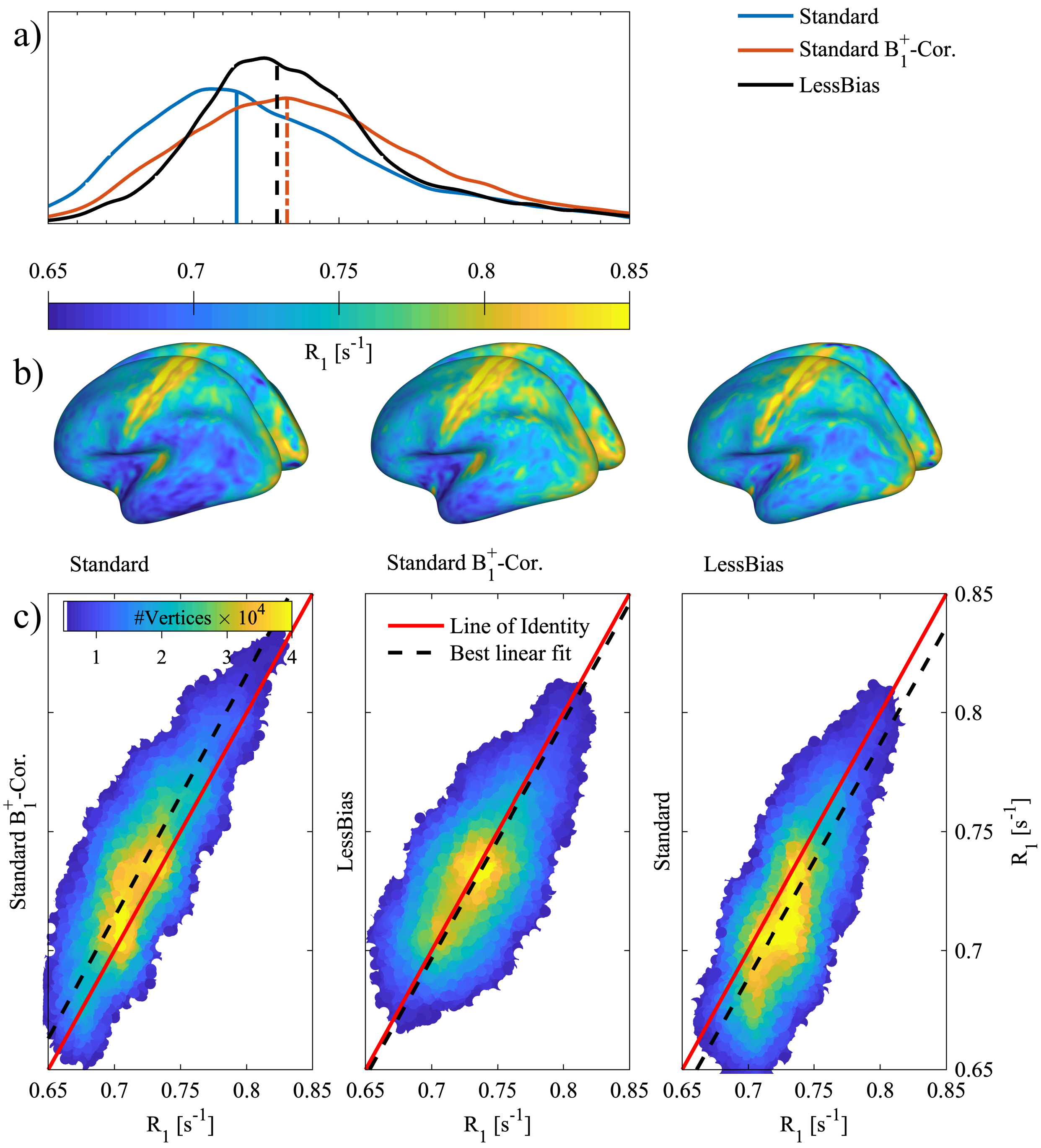


**Figure S13:** GM $R_1$-assessment, Subject 2 Session 1, regarding $B_1^+$-correction. Panel a) shows the distribution of $R_1$-values on the central GM-surface for both Standard and LessBias, As well as for a $B_1^+$-corrected version of Standard. From the distributions and their median values (indicated with vertical lines) it is clear that the $R_1$-values from Standard resemble those of LessBias, when the $R_1$-map from Standard is $B_1^+$-corrected. Panel b) shows the surfaces corresponding to the curves in panel a). A shift towards higher $R_1$-values at the cortex is seen, when the $R_1$-map from Standard is $B_1^+$-corrected. Panel c) shows 2D histograms for the three possible combinations of the maps in panel b). The red lines indicate the perfect relationship between the data and the dashed, black lines indicate the best linear fit with an intercept of zero. It is seen from the plots that while the maps from LessBias and Standard are fairly similar, the best correspondence is found between the $R_1$-maps from $B_1^+$-corrected Standard and LessBias.

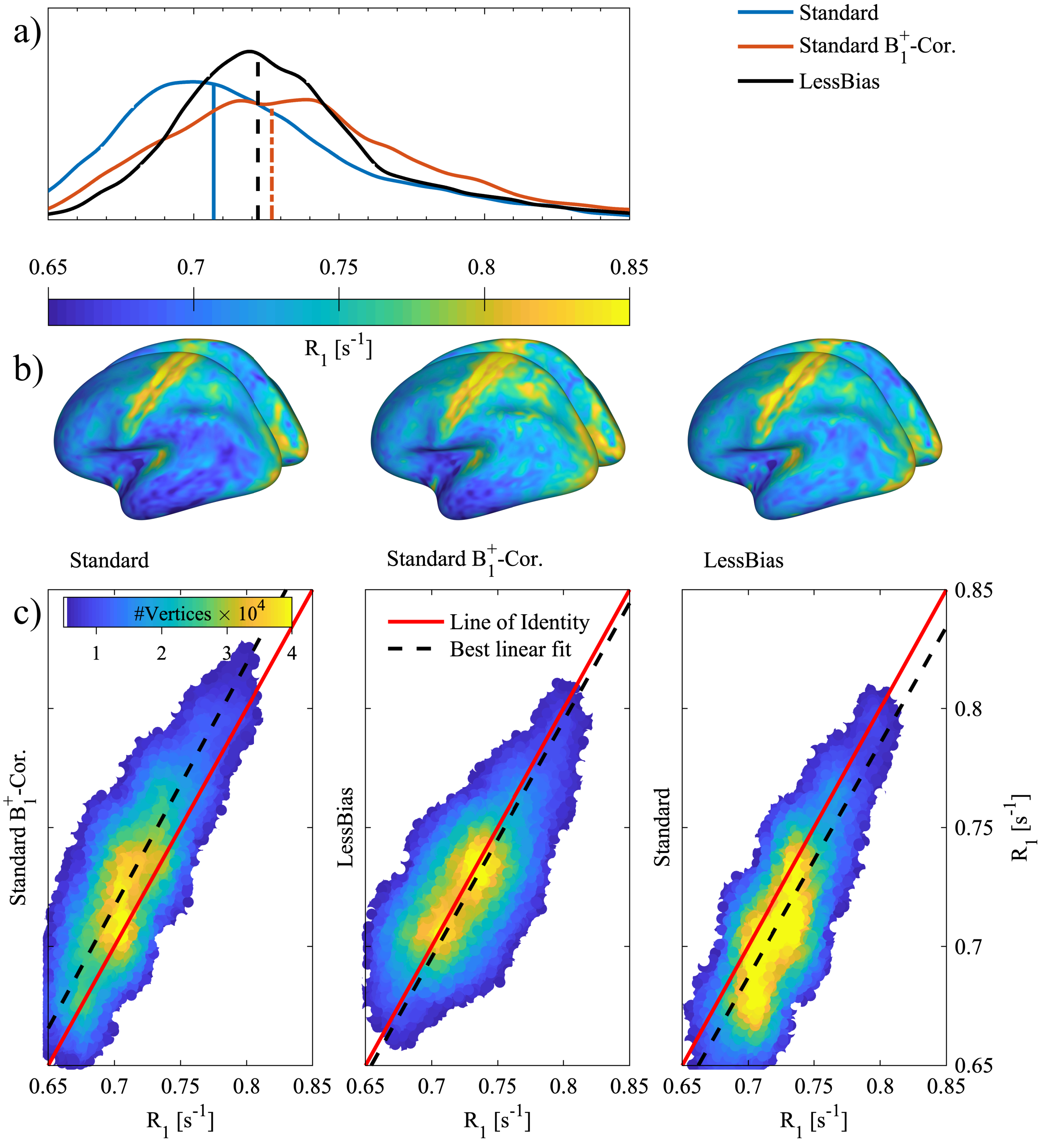


**Figure S14:** GM $R_1$-assessment, Subject 2 Session 2, regarding $B_1^+$-correction. Panel a) shows the distribution of $R_1$-values on the central GM-surface for both Standard and LessBias, As well as for a $B_1^+$-corrected version of Standard. From the distributions and their median values (indicated with vertical lines) it is clear that the $R_1$-values from Standard resemble those of LessBias, when the $R_1$-map from Standard is $B_1^+$-corrected. Panel b) shows the surfaces corresponding to the curves in panel a). A shift towards higher $R_1$-values at the cortex is seen, when the $R_1$-map from Standard is $B_1^+$-corrected. Panel c) shows 2D histograms for the three possible combinations of the maps in panel b). The red lines indicate the perfect relationship between the data and the dashed, black lines indicate the best linear fit with an intercept of zero. It is seen from the plots that while the maps from LessBias and Standard are fairly similar, the best correspondence is found between the $R_1$-maps from $B_1^+$-corrected Standard and LessBias.

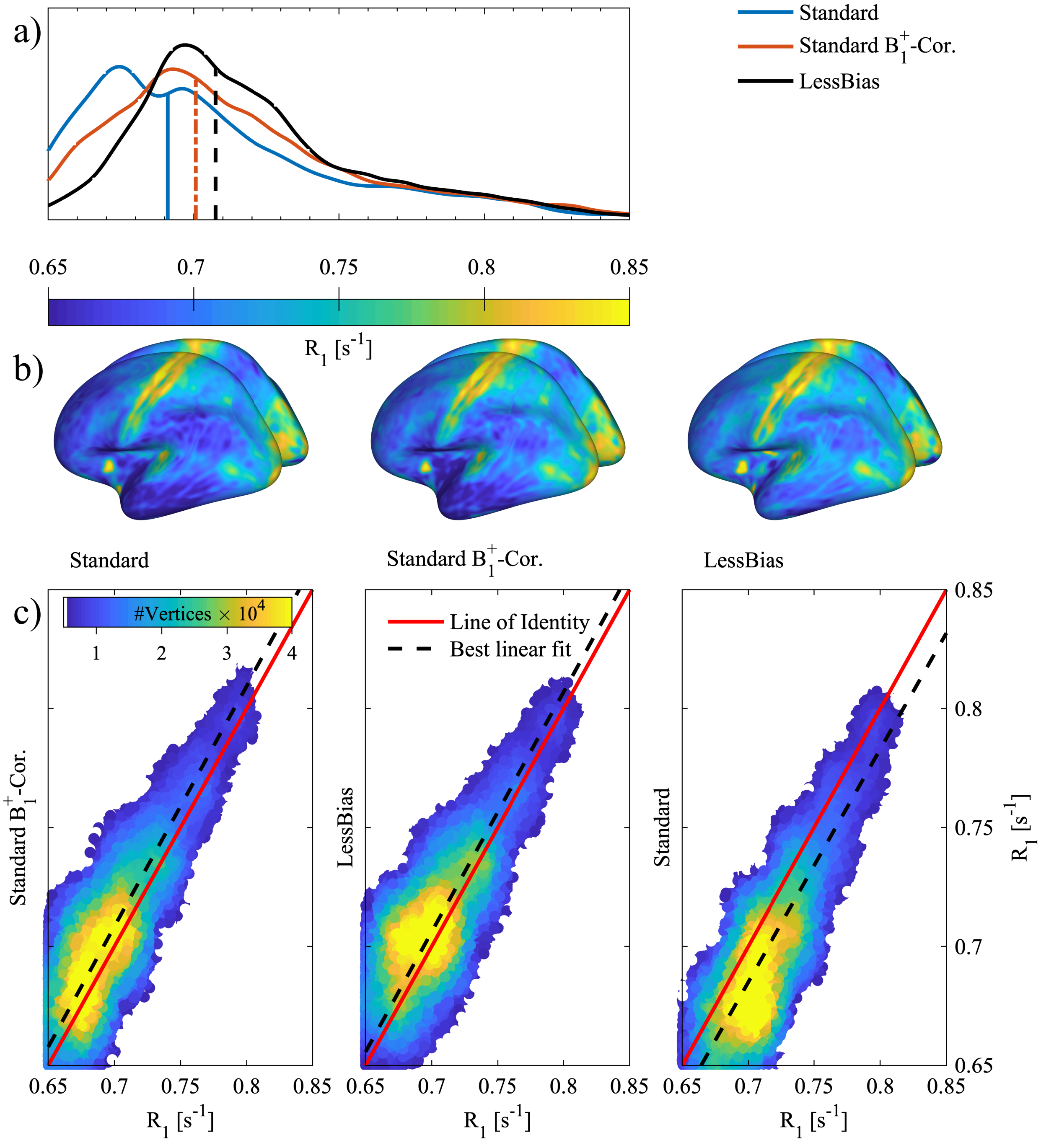


**Figure S15:** GM $R_1$-assessment, Subject 3 Session 1, regarding $B_1^+$-correction. Panel a) shows the distribution of $R_1$-values on the central GM-surface for both Standard and LessBias, As well as for a $B_1^+$-corrected version of Standard. From the distributions and their median values (indicated with vertical lines) it is clear that the $R_1$-values from Standard resemble those of LessBias, when the $R_1$-map from Standard is $B_1^+$-corrected. Panel b) shows the surfaces corresponding to the curves in panel a). A shift towards higher $R_1$-values at the cortex is seen, when the $R_1$-map from Standard is $B_1^+$-corrected. Panel c) shows 2D histograms for the three possible combinations of the maps in panel b). The red lines indicate the perfect relationship between the data and the dashed, black lines indicate the best linear fit with an intercept of zero. It is seen from the plots that while the maps from LessBias and Standard are fairly similar, the best correspondence is found between the $R_1$-maps from $B_1^+$-corrected Standard and LessBias.

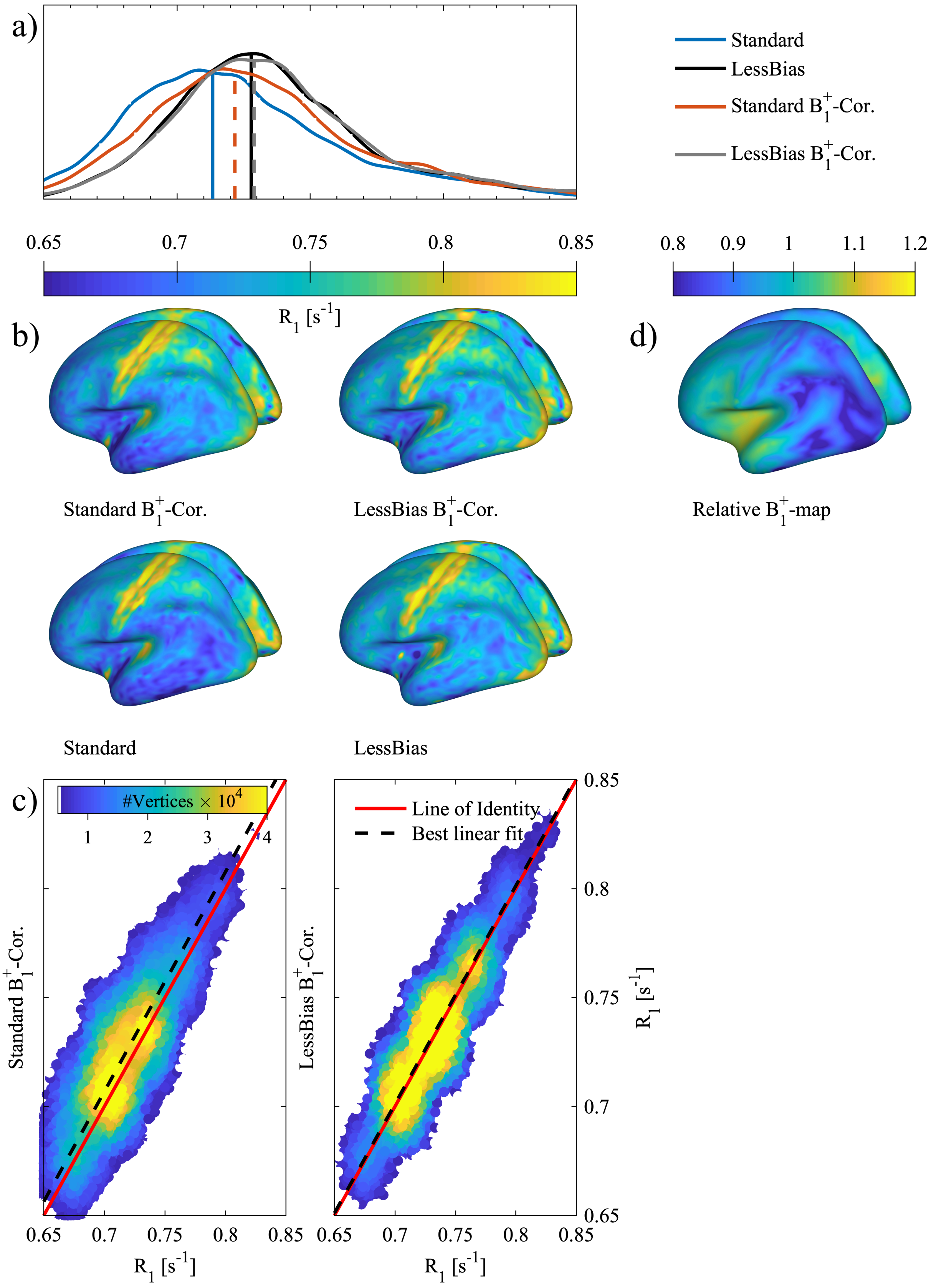


**Figure S16:** GM $R_1$-assessment, Subject 2 Session 1, regarding $B_1^+$-correction. Panel a) shows distributions of $R_1$-values on the central GM-surface for both protocols, with and without $B_1^+$-correction. From the distributions and their median values (indicated with vertical lines) it is clear that the $R_1$-values from Standard resemble those of LessBias, when the $R_1$-map from Standard is $B_1^+$-corrected. However, as could be expected from Figure 5, the effect of $B_1^+$-correction on LessBias is minimal. Panel b) shows the surfaces corresponding to the curves in panel a). A shift towards higher $R_1$-values at the cortex is seen, when the $R_1$-map from Standard is $B_1^+$-corrected. In panel c), 2D histograms show the effect of $B_1^+$-correction for the two different protocols. The red lines indicate the perfect relationship between the data and the dashed, black lines indicate the best linear fit with an intercept of zero. It is seen that LessBias and LessBias with $B_1^+$-correction have a stronger correlation than Standard and Standard with $B_1^+$-correction. Panel d) shows the $B_1^+$-map used for the correction.

### 8 A 7T example with 512 slices

As the desired resolution increases, a larger number of slices is required. Unfortunately this increases the $B_1^+$-inhomogeniety sensitivity, which counteracts the resolution gains if left untreated. Some of this inhomogeneity sensitivity can be counteracted by longer $T_{\mathrm{R,MP2RAGE}}$ and shorter echo spacing, i.e. $T_{\mathrm{R,FLASH}}$.

Additionally, pTx, e.g., with universal pulses, can be used to reduce the severity of the $B_1^+$-inhomogeneity effect, typically to a near 3T like scenario, with a relative $B_1^+ = \pm 20\%$. In Figure S17, we show the transfer function for a Standard* and a LessBias* protocol which have been modified to include 512 slices. Both protocols use $T_{\mathrm{R,MP2RAGE}} = 6\,\mathrm{s}$ and $T_{\mathrm{R,FLASH}} = 3.9\,\mathrm{ms}$. Other parameters were; Standard*: $\mathrm{FA}_1/\mathrm{FA}_2 = 4°/5°$, $T_{\mathrm{I,1}}/T_{\mathrm{I,2}} = 700\,\mathrm{ms}/2500\,\mathrm{ms}$, LessBias*: $\mathrm{FA}_1/\mathrm{FA}_2 = 3°/5°$ , $T_{\mathrm{I,1}}/T_{\mathrm{I,2}} = 500\,\mathrm{ms}/2000\,\mathrm{ms}$. Similar to our 3T example, Figure S17 panel **a** shows that, with a bijective transfer function, a protocol with 512 slices will show substantial $B_1^+$-inhomogeneity sensitivity compared to a protocol with a non-bijective transfer function (Figure S17 panel **b**). One thing to note in Figure S17 panel **b**, is the steeper (negative) slope around the $T_1$ of GM, caused by the higher $T_1$ at 7T compared to 3T. This could lead to slightly increased noise levels in GM, but can be counteracted by an increased $T_{\mathrm{R,MP2RAGE}}$, $T_{\mathrm{I,1}}$ and $T_{\mathrm{I,2}}$ if deemed problematic.

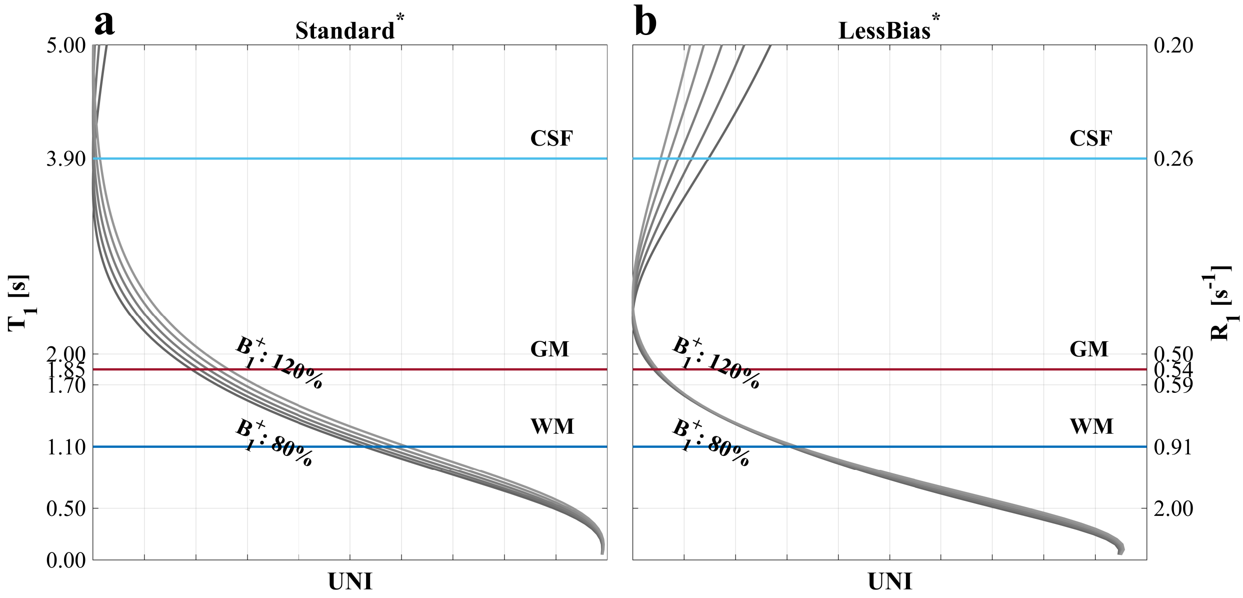


**Figure S17:** Transfer functions for two protocols **a**: Standard* and **b**: LessBias* modified to acquire 512 slices at 7T.
